# Antimicrobial Combination Effects at Sub-inhibitory Doses do not Reliably Predict Effects at Inhibitory Concentrations

**DOI:** 10.64898/2026.02.07.703730

**Authors:** Malte Muetter, Daniel Angst, Roland Regoes, Sebastian Bonhoeffer

**Author notes:** (MM); (SB).

## Abstract

Assessing whether drug combinations synergise or antagonise is difficult for several reasons: (i) measuring bacterial death rates at clinically relevant inhibitory drug concentrations is methodologically challenging, (ii) there is no unifying definition of what constitutes synergy or antagonism, and (iii) both synergy and antagonism may be concentration- and mixing-ratio-dependent. To assess how well sub-inhibitory measurements predict inhibitory behaviour, we quantified drug interactions for 15 pairwise drug combinations on a concentration checkerboard covering a wide range of inhibitory and sub-inhibitory concentrations. To this end, we tracked the population dynamics of 8640 bioluminescent *E. coli* cultures by recording their light-intensity trajectories. To handle time-varying treatment effects and allow fair comparisons between drugs with distinct killing dynamics, we used a time-weighted net growth rate *ψ* to summarise each trajectory and assigned interaction labels (synergistic/independent/antagonistic) based on Bliss independence and Loewe additivity. We found that the interaction label depends on both the concentration and the mixing ratio, frequently changing between the sub-inhibitory and inhibitory regimes. Characterising drug combinations at a single sub-inhibitory concentration is therefore not sufficient. Instead, their combined effects should be assessed at the conditions of their intended use.

## Introduction

Antimicrobial resistance (AMR) poses a major global health burden, with an estimated 1.14 million deaths directly attributable to resistant infections in 2021 alone [1]. The way we use the drugs available to us — what we call the treatment strategy — influences the development and spread of AMR. Often discussed strategies are mixing (randomly assigning drugs to patients), cycling (using only one drug for a certain amount of time, then switching), and combination therapy (each patient receiving multiple drugs simultaneously). Among these, combination therapy appears particularly promising: it is long established against notoriously fast-evolving pathogens such as HIV, malaria, and tuberculosis [2, 3], it outperforms alternatives such as cycling or mixing in theoretical models [4–6], and it performs best in *in vitro* simulations of epidemics [7, 8]. However, clinical evidence for its efficacy is inconclusive, and a comprehensive meta-analysis did not identify a significant overall benefit [9].

Several factors limit what clinical studies can establish about the efficacy of combination therapy, from ethical constraints that exclude the pathogens for which it is already proven beneficial (HIV and *M. tuberculosis*), to a lack of statistical power to track resistance evolution [9]. A further factor is that pooling across combinations may obscure potential differences in efficacy that arise from drug interactions. Drug combinations can exhibit synergistic or antagonistic interactions, depending on whether the combined effect is stronger or weaker than the expectation based on the effects of the single drugs.

Beyond their immediate effect on treatment efficacy, drug interactions also modulate the evolution of resistance. On the one hand, synergy increases the clearance of the pathogen population, leaving fewer opportunities for resistance to develop [10]. On the other hand, synergy tends to select more strongly for resistance than antagonism [10–12], while suppressive antagonism (where the combined effect lies below one of the single-drug effects) can even select against single-drug resistance [13].

Synergy and antagonism are separated by the expected combined treatment effect, which is defined by a reference model. Several such models exist, and which of them is most appropriate to describe independent drugs remains debated [14, 15]. Here, we use two classical and widely applied reference models: the response-based Bliss independence [16] and the dose-based Loewe additivity [17].

Bliss independence is based on the assumption that the probability of being affected by one drug is statistically independent of being affected by the other [16]. This makes Bliss independence more appropriate for drug pairs with distinct and non-overlapping mechanisms of action [18].

In contrast, Loewe additivity – first discussed as iso-additivity by Frei in 1913 [19] and later popularised by Loewe and Muischnek [17] – assumes that the two drugs’ effect-equivalent doses are mutually substitutable.

The expectation from either model is then compared to experimentally observed combined treatment effects. These combined effects are predominantly measured at concentrations below the minimum inhibitory concentration (MIC) (e.g. [20–28]) and at the MIC level (e.g. [29–31]). By contrast, measurements above the MIC, which are clinically and evolutionarily highly relevant [32], remain scarce and often cover only a few points in the inhibitory range (e.g. [33–38]). The primary reason for this is methodological: at sub-inhibitory concentrations, growth rates can be easily assessed at high throughput using optical density measurements, whereas at inhibitory concentrations this typically requires labour-intensive, low-throughput colony-forming unit assays [39]. Interactions are therefore mostly characterised where they are cheapest to measure, and the resulting labels are often portrayed as a property of the drug pair instead of a property of the condition. However, whether interactions measured at sub-inhibitory concentrations align with those at inhibitory ones is questionable, as some studies suggest that interaction labels may depend on dose [21, 40, 41] and on mixing ratio [42].

To systematically test how sensitive the interaction label is to dose and mixing ratio, we measured time-resolved bioluminescence trajectories for all 15 pairwise combinations of six drugs across a wide range of inhibitory and sub-inhibitory concentrations and mixing ratios. Using this dataset, we quantify how often interaction labels based on Bliss independence and Loewe additivity generalise across concentration ranges and mixing ratios.

## Results

We quantified drug interactions for 15 pairwise combinations of six antibiotics falling into three mechanistic classes: polymyxins (colistin, COL; polymyxin B, POL), *β*-lactams (amoxicillin, AMO; penicillin, PEN), and ribosome-targeting drugs (chloramphenicol, CHL; tetracycline, TET). For each drug pair, we assessed 144 conditions (*c*_*A*_, *c*_*B*_) in a 12 *×* 12 checkerboard, with four replicate cultures per condition. For every culture we recorded a light intensity trajectory *I*(*t*) of a bioluminescent *E. coli* MG1655 reporter strain as a proxy for population size.

### Time-variant growth rates and treatment effects

Most drug-interaction metrics are based on estimates of *treatment effects*, which condense the trajectory of the bacterial population size into a scalar. These estimates are typically either rate-based, obtained by fitting an exponential rate of change to a growth trajectory [20, 35, 41, 43, 44], or integral-based, obtained by integrating a (transformed) growth trajectory over time [27, 34, 36, 37, 45, 46]. Rate-based approaches can be used directly to parameterise epidemiological models, and can be estimated in numerous ways, for example from the maximal rate of change or from the net change between endpoints. When a trajectory changes at a constant rate, these methods largely agree and the resulting rate is straightforward to interpret. Under time-varying effects, however, they often disagree and some become unstable. For example, methods based on the minimal or maximal sliding-window fit are sensitive to the shape of the trajectory or to small changes in its endpoint. Integral-based approaches are less exposed to this, as the area under a curve is unambiguous. Both approaches depend on the chosen time window [*t*_1_, *t*_2_].

To allow a fair comparison across drugs with distinct kinetics and durations of action, we use the same window [0, *T*], with *T* ≈ 2 h, for all trajectories (Methods). Given that many trajectories exhibited time-varying treatment effects (see Fig 1a), we summarised each trajectory by *ψ* (Methods), a time-weighted average of the instantaneous net growth rate 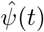 (Equation S2). This weighting emphasises early treatment effects, reflecting that early changes in net growth shape the population for a longer duration than equally sized changes occurring later. *ψ* is proportional to the area under the log-normalised luminescence trajectory *Y* (*t*) = ln(*I*(*t*)*/I*(0)) (Equation S3), and the treatment effect *τ* is proportional to the area between the log-normalised control and treated trajectories (Fig 1b).

**Fig 1.**
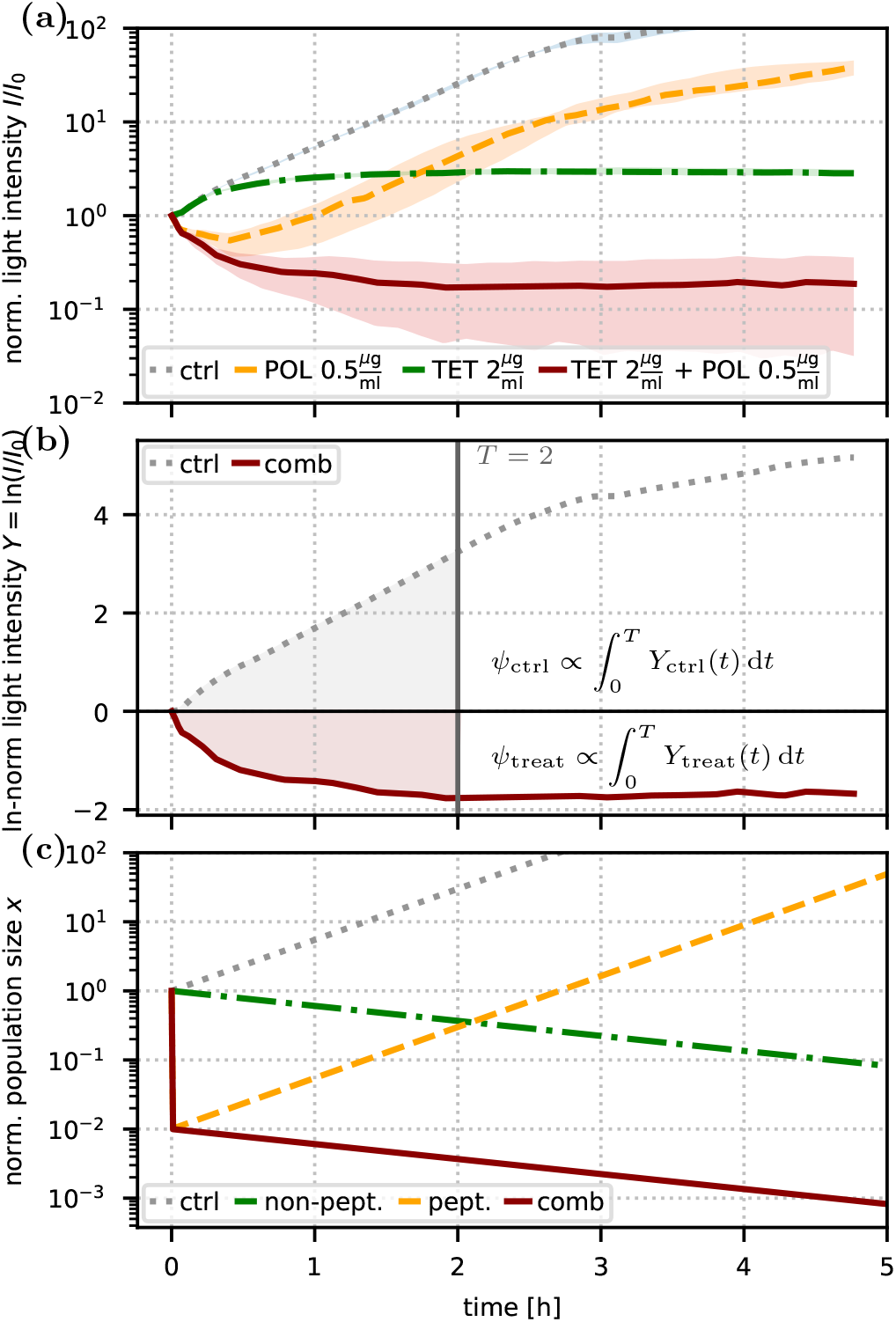
Panel (a) shows the median of normalised light-intensity trajectories *I*(*t*)*/I*(0) for the combination of polymyxin B (POL, 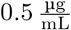) and tetracycline (TET, 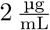), together with the corresponding single-drug and untreated control trajectories, with shaded bands indicating the interquartile range across biological replicates. Panel (b) shows the log-transformed signal *Y* (*t*) = ln(*I*(*t*)*/I*(0)) for the untreated control and the combination treatment from panel (a), together with the corresponding integrals of *Y* (*t*), shaded up to the integration horizon *T* (solid grey line). Panel (c) shows a simple mathematical model illustrating qualitatively distinct dynamics under: no drugs, a continuously acting antibiotic (slow constant decline), a short-acting antibiotic (rapid initial decline, e.g. an antimicrobial peptide), and their combination (rapid initial decline followed by a constant slow decline).

For each drug pair we estimate *ψ* for all checkerboard conditions (*c*_*A*_, *c*_*B*_) jointly (Methods, SI 1G), so that a single *draw* from the estimated distribution holds one *ψ* value for every condition. The resulting time-weighted net growth rates are shown for all combinations in Fig S3.

When two drugs act over distinct time windows, it is not obvious what combined effect to expect. Our dataset contains numerous examples of combinations pairing a peptide, which reduces population size steeply but over a short timeframe, with a second, continuously acting drug. In such cases, the early phase is dominated by the short-acting drug, and the later phase by the continuously acting drug. In Fig 1a, we show a well-behaved example of such a constellation for the peptide polymyxin B (POL), which stops acting after a few minutes, combined with the slower killing but continuously acting tetracycline (TET).

To formally address the question of what treatment effect to expect, we constructed a mathematical model for an idealised scenario. Drug A immediately reduces the population by a factor *α* but does not affect the subsequent dynamics, while drug B acts invariantly over time by constantly reducing the net growth rate from 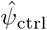 to 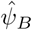. In combination, the population is first reduced by *α* and then grows with 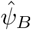 (Fig 1c). Substituting our rate definition (Equation 1) into this model yields additive treatment effects, *τ*_*AB*_ = *τ*_*A*_ + *τ*_*B*_ (see SI 1F).

### Condition-wise interaction score (*µ, ν*) analysis

While the Loewe framework works with any effect definition, for Bliss we first needed to convert the probability-based definition (Equation 3) into a rate-based one, yielding *τ* ^Bliss^(*c*_*A*_, *c*_*B*_) = *τ* (*c*_*A*_) + *τ* (*c*_*B*_) (SI 1D). Notably, the mathematical model above arrives at the same additive expectation, starting from different assumptions.

The interaction scores then measure how far a combination’s treatment effect departs from a reference model’s prediction: the Bliss score *µ* as the normalised deviation from this prediction (Equation 4), and the Loewe score *ν* as the deviation from dose equivalence (Equation 6). Evaluating *ν* additionally requires the single-drug pharmacodynamic curves *f* (*c*), which map a drug’s concentration to the growth rate it produces. Here, *f* (*c*) gives the time-weighted net growth rate *ψ* at concentration *c*. We estimated *f* per drug using a hierarchical (partially pooled) model (Methods, SI 1C; Fig S1, Fig S2). Both scores are evaluated for every eligible condition of every checkerboard, yielding one interaction map per drug pair and reference model (Fig S4, Fig S5). In Fig 2 we show one example checkerboard (CHL+TET) with the estimated *ψ* for every condition (Fig 2a) and the corresponding Bliss interaction map (Fig 2b), illustrating how the interaction changes with concentration (Fig 2c, here from synergy at lower to antagonism at higher concentrations).

**Fig 2.**
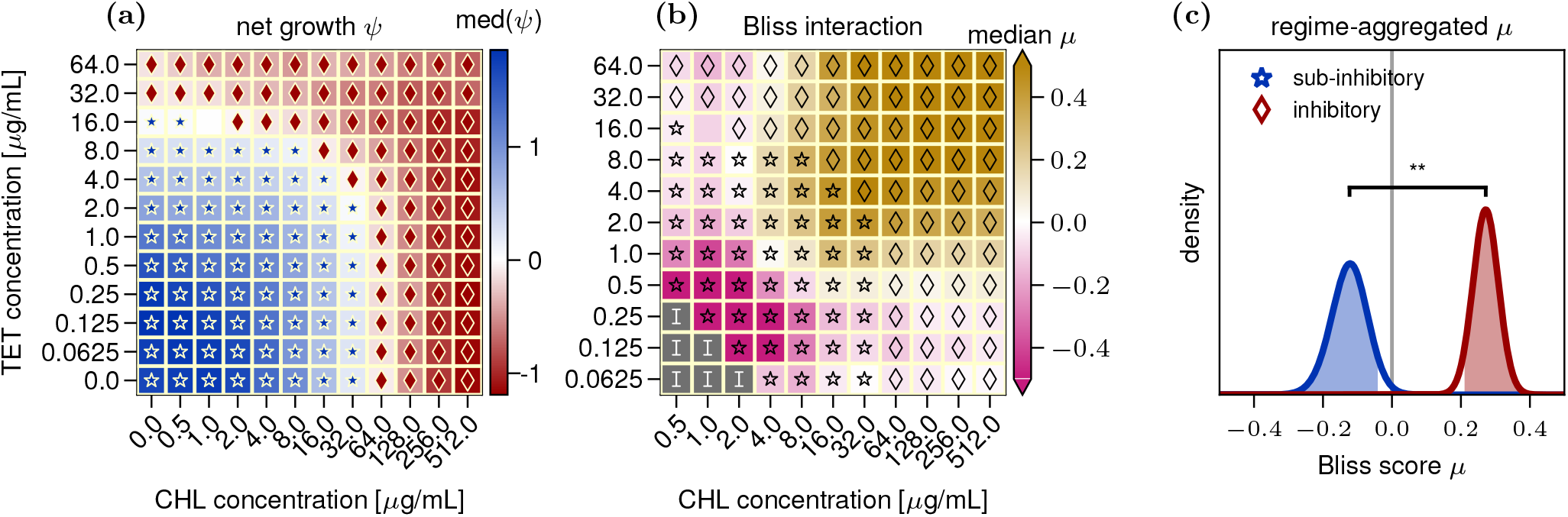
A worked example for the drug combination chloramphenicol (CHL) and tetracycline (TET). **(a)** Time-weighted net growth rate *ψ*: each cell is one concentration pair, coloured by the median *ψ* across *n* = 4 biological replicates. Markers denote regime membership: stars mark sub-inhibitory conditions (net growth), diamonds inhibitory conditions (net killing), assigned when at least 95% of the condition’s *ψ* distribution places it in that regime; unmarked conditions contribute to both regimes (Methods, SI 1G). **(b)** Corresponding heatmap for Bliss independence, coloured by the median interaction score *µ*: magenta indicates synergy (*µ <* 0), gold antagonism (*µ >* 0), and white agreement with the Bliss prediction (*µ* ≈ 0). Markers show the regime membership from (a). Excluded conditions are shown in dark grey, labelled with a code for the exclusion reason; here they all carry “I”, indicating an insignificant treatment effect (Methods). **(c)** Distribution of the regime-aggregated Bliss interaction score *µ* for the inhibitory (red) and sub-inhibitory (blue) regimes (Methods, SI 1G). A regime is classified as synergistic (*µ <* 0) or antagonistic (*µ >* 0) when at least 95% of that distribution (the shaded area under the curve) lies on that side of zero, and independent otherwise. When the two regimes carry opposite labels, we call this a strong disagreement. The bracket compares the regime scores: **\*** if at least 95% of their difference lies on one side of zero, **\*\*** if at least 99% does. Here, the sub-inhibitory regime is inferred to be synergistic and the inhibitory antagonistic (strong disagreement).

### Interaction scores often differ between concentration regimes

To assess whether interactions remain consistent across concentration regimes, we first assign each eligible condition to the inhibitory or sub-inhibitory regime with conditions spanning both regimes contributing to both (Methods). We then aggregate the condition score distributions regime-wise via a Bayesian random-effects meta-analysis (Methods, SI 1G), yielding one score distribution per regime (Fig 2c). Finally, we test whether the two regimes differ significantly, i.e. whether at least 95% of the difference between their distributions lies on one side of zero (SI 1G). This was the case for 7 of 15 pairs under Bliss and 8 of 15 under Loewe (Fig S6, Fig S7).

### Inferred interaction labels often disagree between concentration regimes

We then assign a regime-wide interaction label: antagonism if at least 95% of the score distribution lies above zero, synergy if at least 95% lies below zero, and independence otherwise (Methods). The resulting medians and labels are reported in Fig 3a,b and Table S2. To compare regimes, we define three alignment classes: agreement (same label), soft disagreement (independent in one but synergistic or antagonistic in the other), and strong disagreement (opposite labels).

**Fig 3.**
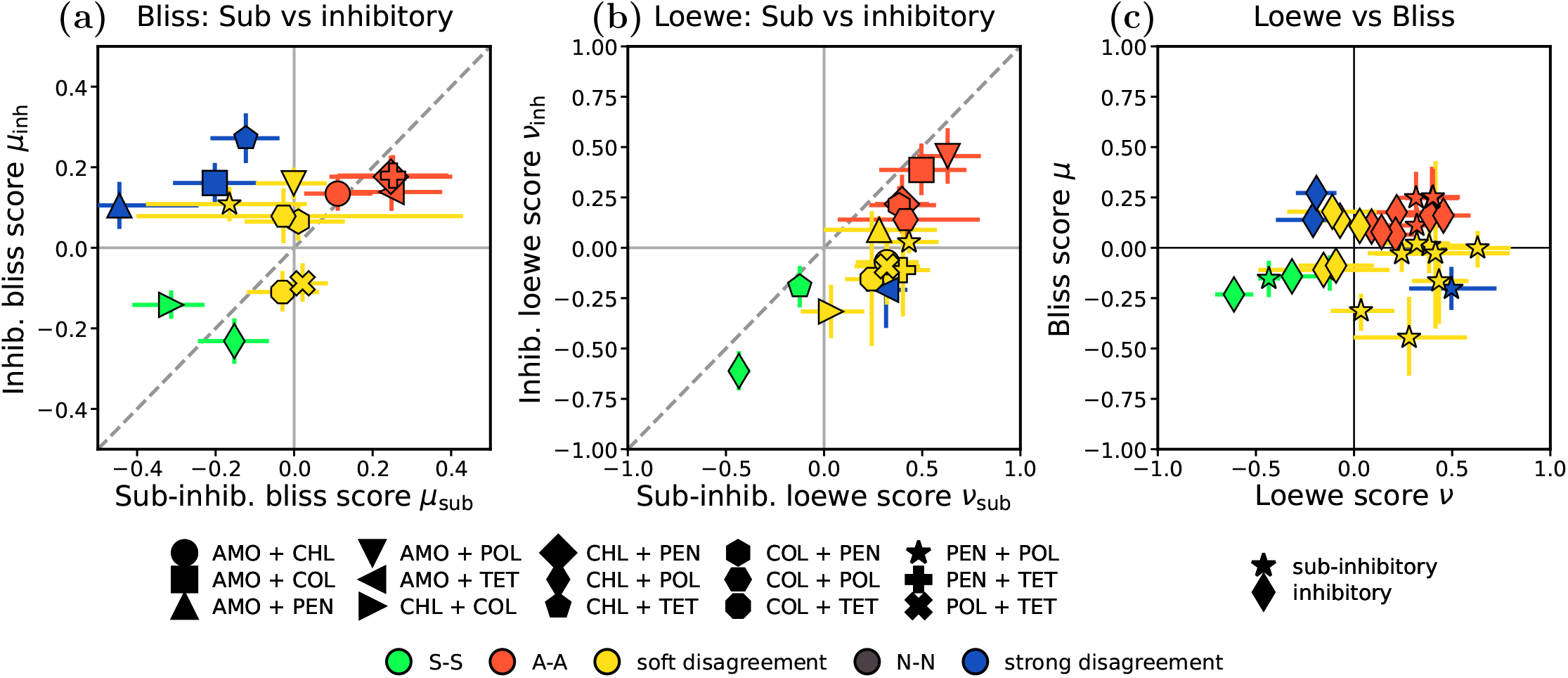
Interaction summaries across regimes and models. Panels **(a)** and **(b)** compare sub–inhibitory vs. inhibitory interaction scores, using **(a)** the Bliss interaction score *µ* and **(b)** the Loewe interaction score *ν*. Each point represents one drug combination, plotted as the median estimate in the sub–inhibitory regime (x-axis) against the median estimate in the inhibitory regime (y-axis), with error bars showing the central 90% interval. In panels **(a)** and **(b)**, markers encode the drug combination from one shared legend, and the dashed diagonal marks *y* = *x* (equal sub–inhibitory and inhibitory score), i.e. where the concentration regime does not change the score. **(c)** Loewe vs. Bliss comparison across regimes: each point corresponds to one combination and regime, plotted as the median Loewe interaction score *ν* (x-axis) against the corresponding Bliss interaction score *µ* (y-axis), with error bars showing the central 90% interval and marker shape encoding the regime. Across all panels, we classify interactions as synergistic (S), antagonistic (A), or independent (N; not significantly different from zero); a regime is independent exactly when its error bar covers zero. Colours indicate interaction label agreement between the two compared axes in each panel: *agreement* if the two interaction labels coincide (N–N, S–S, A–A), *soft disagreement* if one interaction label is independent and the other is non-independent (N–S, N–A), and *strong disagreement* if the classifications are opposite (S–A).

Assuming Bliss independence, we observed agreement for six combinations and disagreement for nine, of which three were strong (AMO+COL, AMO+PEN, CHL+TET) and six soft. Assuming Loewe additivity, we observed agreement for seven combinations and disagreement for eight, of which one was strong (AMO+TET) and seven soft.

The two comparisons identify overlapping but distinct sets of pairs: under Loewe, seven of the eight label disagreements were accompanied by a significant difference between the regime scores; under Bliss, six of nine. Conversely, under each reference model, one pair carried the same label in both regimes although its scores differed significantly.

### Interaction labels are sensitive to analysis choices

To assess the robustness of the regime labels, we repeated the analysis under eight alternative configurations, varying the rate estimator, the aggregation, the time window and the sampling of mixing angles (Methods, SI 1H; Fig S8, Fig S9, Fig S10).

Relative to the base analysis, every configuration changed at least 6 of the 60 regime labels (15 pairs *×* 2 regimes *×* 2 reference models). The largest change came from replacing the meta-analysis with pooling every eligible condition’s draws into a single sample. This widens the intervals roughly sixfold, so that all but the most extreme regimes are classified as independent, changing 39 labels. Importantly, every configuration still produced pairs whose regime labels disagree.

### Interaction labels depend on the reference model

Fig 3c replots the same interaction summaries as in panels (a) and (b), but now compares the Bliss interaction score *µ* to the Loewe interaction score *ν* across both sub-inhibitory and inhibitory regimes. Across all 30 (2 *×* 15) comparisons, the Bliss- and Loewe-based classifications agree in 14 cases, show soft disagreement in 13, and strong disagreement in three (AMO+COL, AMO+TET, CHL+TET). Regime agreement is likewise model-dependent: six pairs carried the same label in both regimes under Bliss and seven under Loewe, but only two did so under both reference models.

### Interaction labels can change with dose even at fixed mixing ratio

Above, we compared interaction summaries between sub-inhibitory and inhibitory regimes by aggregating condition-wise estimates across a range of doses and mixing ratios. We next ask whether interaction labels also change (i) as the dose increases at a fixed mixing ratio, and (ii) as the mixing ratio varies at a fixed effect level. For both analyses, we normalise each drug’s concentration by its fitted zMIC, the concentration at which *ψ* = 0 (Equation 2). We then reparameterise each pair of normalised concentrations in polar coordinates (*z, φ*), where the radius *z* is the *combined dose* and the angle *φ* is the *mixing angle*, which encodes the ratio of the two normalised concentrations (Equation 7).

For each drug combination, we build a continuous, monotone consensus surface *ψ*^cons^(*c*_*A*_, *c*_*B*_) from the estimated interaction field (SI 1J; Fig S11). From this surface, we estimate *polar pharmacodynamic curves*, which show the net growth rate *ψ* as a function of the combined dose *z* at a fixed mixing angle *φ* = 45^*◦*^ (equal normalised concentrations). At a given dose, predicted values above the observed *ψ* indicate synergy, and values below it antagonism. For both reference models, there are examples where the interaction changes direction as the combined dose increases. For the combination AMO+COL (Fig 4a), the Loewe-based interaction shifts from antagonism at lower doses to synergy at higher doses. For AMO+PEN (Fig 4b), the Bliss-based interaction flips, with synergy at lower doses and strong antagonism at higher doses. *Polar pharmacodynamic curves* at 45^*◦*^ for all combinations are shown in Fig S12.

**Fig 4.**
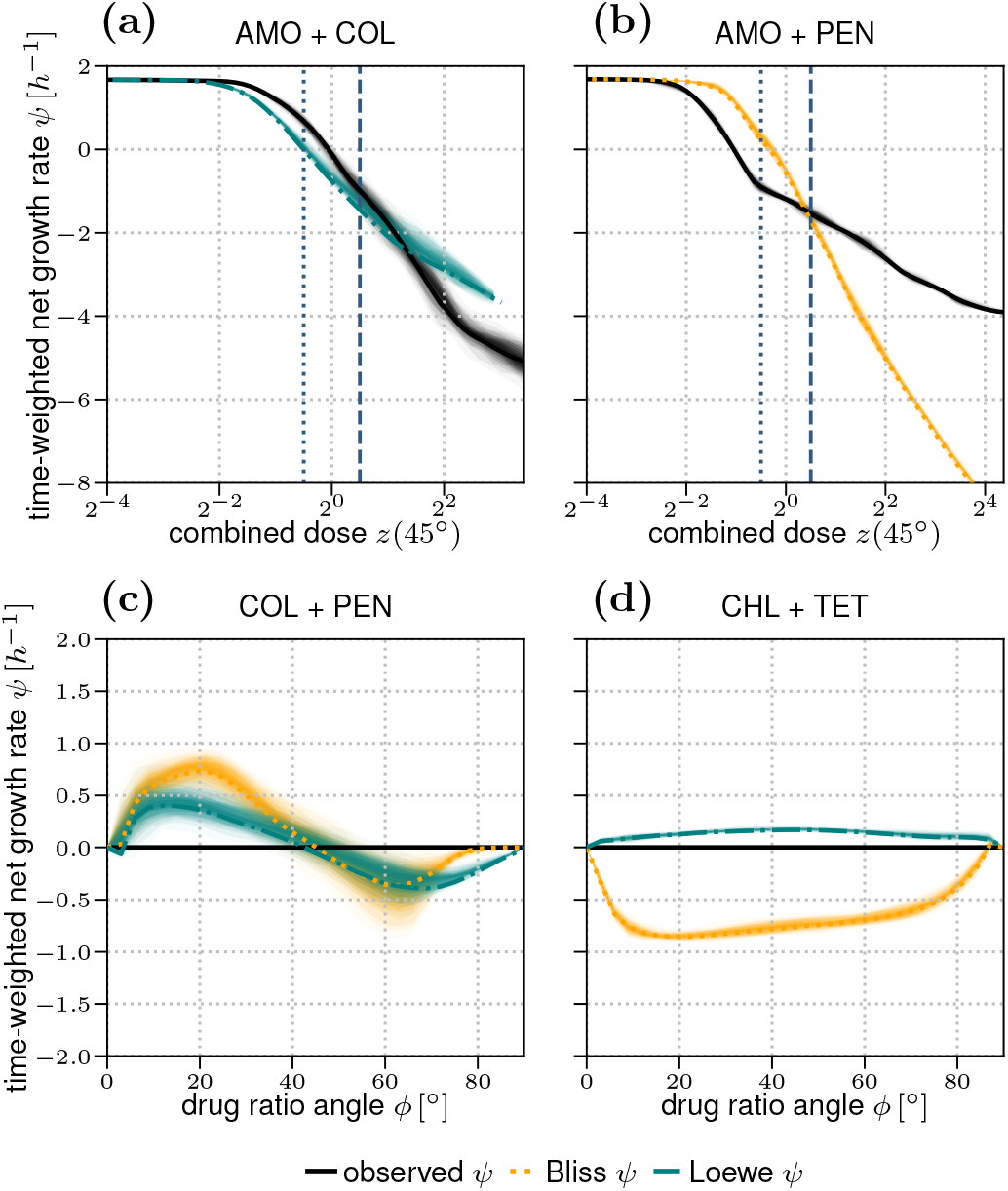
Panels (a,b) show *polar pharmacodynamic curves* at *φ* = 45^*◦*^ (corresponding to a 1:1 ratio in units of zMIC) for (a) AMO+COL and (b) AMO+PEN. The x-axis shows the combined dose *z*, where 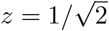 (blue dotted line) corresponds to both single-drug doses equaling 0.5 zMIC, and 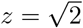 (blue dashed line) corresponds to both equaling 1 zMIC. Panels (c,d) show the Bliss- and Loewe-based predictions for *ψ* over the mixing angle *φ* along the observed isobole at 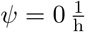 for (c) COL+PEN and (d) CHL+TET. Shaded regions show the uncertainty of the surface estimate (SI 1J).

### Interaction labels can depend on the mixing ratio

To assess whether interaction labels depend on the mixing ratio at a fixed effect level, we extract isoboles from the continuous surface fits (SI 1J; Fig S13). Along each isobole, we evaluate Bliss- and Loewe-based predictions and plot them as a function of *φ*. For many combinations, the inferred interaction label is stable across *φ*, as exemplified by CHL+TET (Fig 4d). However, combinations showing mixing-ratio dependence are also common, such as COL+PEN (Fig 4c). Plots for all combinations are shown in Fig S14.

## Discussion

We quantified interaction patterns across a wide concentration range for several drug combinations with different modes of action. Quantifying the size of the bacterial population is notoriously difficult because bacterial death has many facets, and no single method captures all of them [47, 48]. In this work, we used bioluminescence as a proxy for population size. This choice enabled us to record 8640 finely time-resolved (every 10 minutes for five hours) growth trajectories at high throughput. Bioluminescence has been shown to be a better proxy for biomass than for cell number [48], so our readout is most robustly interpreted as biomass. However, since we restricted our analysis to drugs for which changes in biomass mostly align with changes in cell number (i.e. limited filamentation) [48], we do not expect qualitatively different conclusions when using an alternative readout of population size.

For many of the observed trajectories treated with higher concentrations of peptides (COL and POL), the light intensity dropped below the detection limit within the first few minutes. In principle, rates could still be estimated for such curves over a shorter or shifted window. However, comparing estimates obtained over different windows across drugs with different onsets and durations of treatment effects is conceptually difficult to justify. We therefore imposed the same window on all trajectories (*T* ≈ 2 h). This requires valid data across that window and comes at the cost of discarding all trajectories that fall below the detection limit too early, which means losing more than half of the trajectories for COL and POL. For conditions where only a part of the replicates survived this introduces a bias towards weaker killing, since faster-declining replicates are the most likely to be discarded.

To address our core question — how well sub-MIC measurements predict interactions at inhibitory concentrations — we aggregated interaction estimates for sub-inhibitory and inhibitory conditions under both reference models (Bliss independence and Loewe additivity), and compared the two regimes in two ways: whether they carry the same interaction label, and whether their scores differ. Label disagreements between the two regimes were common. Most of these were soft, i.e. a significant synergy or antagonism in one regime but independence in the other. Soft disagreements are not straightforward to interpret, as an independence label can arise in several ways. The most obvious is that the interaction in that regime genuinely is independent. But it can also indicate a concentration dependence within the regime that averages out to independence, which can happen whenever the boundary between synergy and antagonism does not coincide with the line separating the two regimes (*ψ* = 0). A good example is CHL+TET under Bliss, which is synergistic at lower and antagonistic at higher concentrations (Fig 2b). Our main analysis classifies this pair as a strong disagreement, but four of the alternative configurations label it a soft one (Fig S9i), because the line between the interaction types runs through the sub-inhibitory regime. Further reasons for an independence label include angular dependence (Fig S14), few contributing regime conditions, and experimental noise.

Concentration dependence can also be seen by asking whether the regime scores differ significantly, rather than whether each regime separately clears the threshold. This sidesteps two ways in which the label comparison can mislead. A label can change while the scores stay nearly identical, as for (among others) COL+PEN under Bliss, where it moves from independent to antagonistic between medians of 0.011 and 0.065. Conversely, a label can stay the same while the scores differ substantially, as for CHL+COL under Bliss, where both regimes are called synergistic although their scores differ significantly (medians −0.313 and −0.142).

Our results also confirm the practical limitations of Loewe-based interaction measures at high concentrations that have been reported previously [49]. Since Loewe relies on the inverse of the single-drug pharmacodynamic functions, it is only defined when the combined effect lies within the effect ranges reached by both single-drug treatments. For drug pairs with very different maximal killing rates, this condition fails in large parts of the checkerboard (cells marked L in Fig S5). This limitation prevents the quantification of drug interactions for many therapeutically relevant conditions.

Our findings show that conclusions about synergy or antagonism are fragile in two distinct ways. The first is inherited from the difficulty of quantifying time-variant killing. The choice of rate estimator or time window frequently changed the estimated interaction labels (Methods, SI 1H; Fig S8, Fig S9, Fig S10). The second is inherent to the interaction scores themselves. They depend on the concentration regime, on the mixing ratio and, as previously reported [50], on the reference model (Fig 3, Fig S14). Accordingly, single-point sub-inhibitory measurements are insufficient to reliably characterise drug interactions at clinically relevant inhibitory concentrations, and interaction scores must instead be assessed directly at the conditions of interest.

## Materials and methods

### Strains and Media

We used the bioluminescent strain *Escherichia coli* MG1655 ΔgalK::(kan^*R*^-luxCDABE) constructed previously [48]. Cultures were grown in LB medium (Sigma L3022).

### Drug preparation

We explored drug interactions among six drugs (Table 1), resulting in 15 drug–drug pairs. For each pair, one compound was designated drug A and the other drug B. Each compound was prepared as a 20*×* stock of its highest working concentration. We then performed a twofold dilution series in a 12-column deepwell plate, resulting in 11 diluted concentrations and one drug-free column. For drug A, we transferred 125 µL per well into a 96-well plate (subreservoir A) with a horizontal concentration gradient. For drug B, we transferred 50 µL per well into two 96-well plates (antibiotic reservoir plates I and II) with vertical concentration gradients (each 6 *×* 12 layout). All plates were stored at −80 °C to minimise degradation over time, accepting a single freeze–thaw cycle.

**Table 1.**
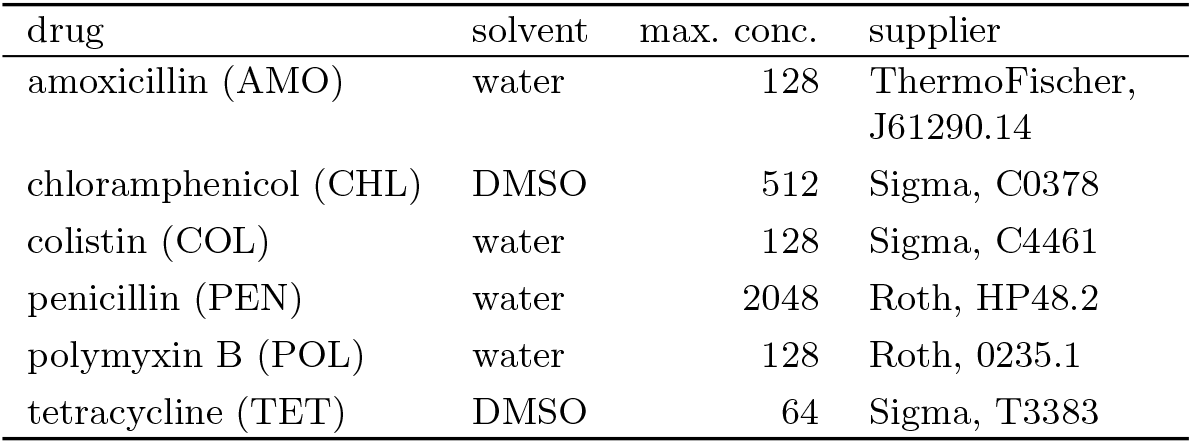
Drugs used in the experiments, the solvent, the highest concentration tested 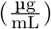, and the supplier with catalogue number. See Table S1 for the estimated zMIC of each drug.

| drug | solvent | max. conc. | supplier |
| --- | --- | --- | --- |
| amoxicillin (AMO) | water | 128 | ThermoFischer, J61290.14 |
| chloramphenicol (CHL) | DMSO | 512 | Sigma, C0378 |
| colistin (COL) | water | 128 | Sigma, C4461 |
| penicillin (PEN) | water | 2048 | Roth, HP48.2 |
| polymyxin B (POL) | water | 128 | Roth, 0235.1 |
| tetracycline (TET) | DMSO | 64 | Sigma, T3383 |

Because each drug is prepared as a single solvent-matched stock and then serially diluted, the solvent concentration scales with the drug concentration. We did not separate drug from solvent effects, but treated them together as solvent–drug hybrids. Our results therefore strictly apply to these hybrids rather than to the pure drugs; for readability we state this caveat once here rather than repeating it throughout. Treated as hybrids, all results are internally consistent, while the treatment effects measured in the highest concentrated wells may exceed those expected using the pure drugs. This is most relevant for CHL and TET, which were dissolved in DMSO, with the single highest concentration of DMSO reaching 10 %. We listed all DMSO concentrations in Table S5 and Table S6, and marked cells with concentrations equal to or exceeding 1.25 % in red.

### Dose response assays

For each assay — one per drug combination — we prepared four overnight cultures grown for 14 h. Cultures were distributed across a white 384-well plate (Greiner 781073) used as the source plate, so that the replicates formed a 2 *×* 2 block format (e.g. C1–rep1, C2–rep2, D1–rep3, D2–rep4). We then filled two assay plates (white 384-well plate) with 54 µL LB per well and transferred inocula from the source plate to both assay plates (I, II) using an Evo 200 liquid handling platform (Tecan) with a pintool. The dilution was approximately 1:150, although we observed substantial well-to-well variation in the effective inoculum size (see below). Plates were incubated for 2 h to reach exponential phase. During this incubation, subreservoir A and both antibiotic-reservoir plates were thawed, and 50 µL from each well of subreservoir A was transferred to the corresponding wells of both antibiotic-reservoir plates, generating a 10*×* mixture of drugs A and B.

For both assay plates, the assay started as follows: a baseline luminescence reading was taken, 6 µL of the 10*×* drug mixture was added (defining *t* = 0 at dosing for the respective wells), and a second luminescence reading was taken. Subsequently, we alternated between reading assay plates I and II for a total duration of 5 h. Luminescence was recorded with an Infinite F200 plate reader (Tecan) using a 250 ms integration time.

### Data preprocessing

To estimate stray-light contamination, we conducted a calibration experiment in which six source wells (E5, E12, E20, L5, L12, L20) contained stationary-phase cultures while all other wells remained empty. From a single luminescence read of the full plate, we constructed a distance-dependent stray-light kernel (Fig S15) and corrected each well by subtracting the summed contributions from neighbouring wells (restricted to distances *d* ≤ 3, Equation S8). We defined the lower detection limit as *I*_LoD_ = 10 RLU and the upper detection limit as 10^6^ RLU. We defined the common analysis horizon *T* as the earliest time point at which any untreated control well exceeded the upper detection limit, yielding *T* ≈ 2 h. For each well, we computed the fraction of observations up to *T* below *I*_LoD_. Wells with more than 20 % of observations below *I*_LoD_ were excluded from all subsequent analyses. For retained wells, values below *I*_LoD_ were censored by replacing them with *I*_LoD_. For numerical integration, we interpolated the log-normalised signal to obtain values at the common time horizon *t* = *T* (see also SI 1B).

### Time-weighted net growth rate

We define *ψ* as the linearly time-weighted net growth rate (SI: Equation S2) over the common time horizon *T*. We infer *ψ* by calculating the scaled integral of the log-normalised light intensity *Y* (*t*) = ln (*I*(*t*)*/I*(0)) (SI: Equation S3),

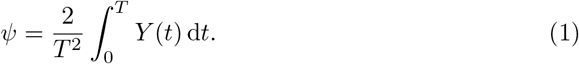

### Single-drug pharmacodynamic curve fitting

For each drug, we modelled the pharmacodynamic curve *f* (*c*) as a Hill function [43]:

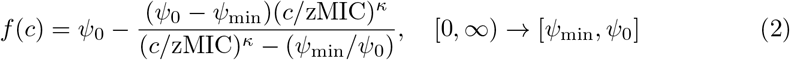

Here, *ψ*_0_ denotes the maximum and *ψ*_min_ the minimum time-weighted growth rate, corresponding to *c* = 0 and *c* → ∞, respectively. *κ* denotes the Hill coefficient and zMIC the concentration at which *f* (zMIC) = 0. *ψ*_min_ is not identifiable from our data, since the concentrations measured do not reveal whether *ψ* would fall further at higher doses. Since *ψ*_min_ must lie below the lowest rate observed for that drug, we fix it 5% below that value. The remaining parameters were estimated hierarchically across the experiments each drug appears in, by Hamiltonian Monte Carlo (NUTS [51]) in PyMC [52] (SI 1C; Table S1). Prior widths were calibrated to the data (Table S3), and we assessed their influence on the posteriors by widening and tightening all priors by a factor of two (SI 1I; Table S4).

### Interaction field

For each drug pair, all checkerboard conditions are estimated jointly as one Gaussian Markov random field [53]. Each condition is informed by its own valid replicates and by its neighbours, through a Gaussian smoothing prior (SI 1G, SI 1I). We fit the field by Gibbs sampling: each sweep updates every condition once and yields one posterior draw of the entire field, within which *τ*, *µ* and *ν* are computed. We run two chains of 2500 sweeps, discarding the first 600 and keeping every second sweep thereafter, giving 1900 retained draws in total. The chains converged 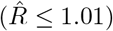, and the retained draws are only weakly autocorrelated, with a bulk effective sample size of 1853 of 1900 at the median and never below 500 [54]. Stacking the draws gives the posterior distributions of *ψ, τ*, *µ* and *ν*.

### Bliss-based interaction score *µ*

Bliss independence assumes that the probabilities of being affected by each drug are statistically independent,

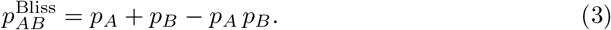

For time-varying hazards, this implies additive treatment effects, *τ*_*AB*_ = *τ*_*A*_ + *τ*_*B*_ (SI 1D, Equation S15). We define the Bliss-based interaction score as the normalised deviation from that prediction:

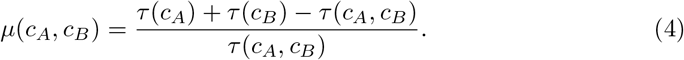

To avoid division by zero, we only evaluate *µ*(*c*_*A*_, *c*_*B*_) if *τ* (*c*_*A*_, *c*_*B*_) is significantly greater than zero (5th percentile of its draws *>* 0).

### Loewe-based interaction score *ν*

Under Loewe additivity, the isobole — the set of concentration pairs producing the same effect *ψ* — is the straight line connecting the equivalent single-drug doses, so that

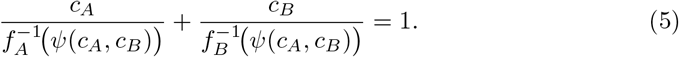

The inverse 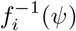 is the corresponding single-drug equivalent concentration of drug *i*. We define a Loewe interaction score (details in SI 1E) as the deviation from this condition:

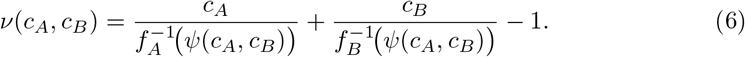

Our definition of *ν* is analogous to the combination index (CI − 1) [40], but uses a different pharmacodynamic model *f*_*i*_ (either Equation S10 or inferred numerically from the single-drug edges of the surfaces, SI 1J). We only evaluate *ν*(*c*_*A*_, *c*_*B*_) if *τ* (*c*_*A*_, *c*_*B*_) is significantly greater than zero. In addition, *ν* is defined only where the combined effect lies within the effect range of both single drugs, *ψ*(*c*_*A*_, *c*_*B*_) ≥ max(*ψ*_min,*A*_, *ψ*_min,*B*_) (Equation S19), which we require for at least 10% of posterior draws.

### Concentration regimes

Conditions are split draw-wise into two concentration regimes by the sign of their net growth rate: sub-inhibitory, where the population still grows (*ψ >* 0), and inhibitory, where it declines (*ψ <* 0). A condition contributes to a regime if at least 5% of the draws place it there, and conditions whose *ψ* distribution straddles zero contribute to both.

### Regime aggregation

Within a regime, each contributing condition *i* is summarised by its median score *y*_*i*_ (i.e. *µ*_*i*_ or *ν*_*i*_) and the standard deviation *s*_*i*_ of that score, both computed from that condition’s in-regime draws only. These are aggregated by a Bayesian random-effects meta-analysis [55], 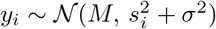, where *M* is the regime-level score and *σ* the spread between conditions (SI 1G). We report *M* ‘s median and central 90% interval; the interval excluding zero is equivalent to at least 95% of the posterior lying on one side of zero.

A regime is labelled antagonistic if that 95% lies above zero, synergistic if it lies below, and independent otherwise. To compare the two regimes directly, we apply the same criterion to the difference *M*_inh_ − *M*_sub_, marking it **\*** at 95% and **\*\*** at 99%.

### Polar reparametrisation

We normalise concentrations of drug *i* using the corresponding zMIC_*i*_ estimates obtained from pharmacodynamic curve fits (Table S1): *z*_*i*_ = *c*_*i*_*/*zMIC_*i*_. We then define polar coordinates based on the normalised concentrations:

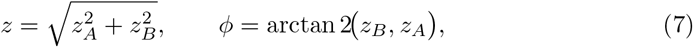

with *φ* corresponding to the *mixing angle* and *z* to the *combined dose*. These coordinates were used to plot polar pharmacodynamic curves, which show the net growth rate *ψ* over the combined dose *z* for a fixed mixing angle *φ*. These curves represent cross-sections through the surfaces at fixed *φ* (SI 1J).

### Inoculum effects

We observed a larger-than-expected variation in pre-treatment light intensity *I*_0_, which we use as a proxy for inoculum size. Regressing *ψ* on *I*_0_ for each single-drug condition and concentration showed negligible effects for AMO, CHL, PEN, and TET, but substantial effects for COL and POL at intermediate concentrations (Fig S16). Because inoculum size showed no significant directional trend along the concentration index (*p* = 0.092), this variation primarily adds noise and contributes to the increased spread in Fig S2c,e (SI 1K).

## Data, materials and software availability

The experimental data are available at Zenodo (DOI: 10.5281/zenodo.22079689) and the analysis code at Zenodo (DOI: 10.5281/zenodo.18374151). Supplementary methods, figures and tables are provided in S1 Text.

## Supporting information

### S1 Text. Supplementary methods, figures and tables

Derivations of the time-weighted net growth rate, the Bliss and Loewe interaction scores, the hierarchical pharmacodynamic model, the interaction-field estimation and the regime-level aggregation, together with the sensitivity analyses and all supplementary figures and tables.

## Acknowledgments

We thank ETH Zurich for funding this work. During manuscript preparation, we used OpenAI’s ChatGPT and Anthropic’s Claude Opus for editorial assistance (coding, grammar, and proofreading).

## Notes

https://doi.org/10.5281/zenodo.22079690

https://doi.org/10.5281/zenodo.22079412

### Competing Interest Statement

The authors have declared no competing interest.

### Summary of Updates

This version differs from the previous one in wording and presentation; the data, the analysis and the conclusions are unchanged. We simplified the notation of the main text: distributions are no longer set in bold, and the sampler terminology is confined to the Methods, where it is defined. Several passages in the Results and the Discussion were shortened, and the supplementary methods were expanded where they were too terse.

