## supplementary information for "Antimicrobial Combination Effects at Sub-inhibitory Doses do not Reliably Predict Effects at Inhibitory Concentrations"

### SI TEXT

**Notation.** Treatment-dependent quantities $X$ are written as functions of the underlying condition, $X(\{c_i\})$, where $\{c_i\}$ is the set of drug concentrations defining that condition.

**1A. Time-weighted net growth rates.** We define $\psi$ as the linearly time-weighted average of the temporal net growth rate $\hat{\psi}$:

$$w(t) = \frac{2}{T^2}(T-t), \tag{S1}$$

$$\psi = \int_0^T \hat{\psi}(t)\, w(t)\, dt = \frac{2}{T^2}\int_0^T (T-u)\,\hat{\psi}(u)\, du, \tag{S2}$$

where the kernel $w(t)$ decreases linearly over time and is normalised such that $\int_0^T w(t)\,dt = 1$. To derive $\psi$ from the luminescence trajectories $I(t) = I_0 e^{\int_0^t \hat{\psi}(u)\,du}$, we define the log-normalised trajectory

$$Y(t) = \ln\frac{I(t)}{I(0)} = \ln I(t) - \ln I(0) = \int_0^t \hat{\psi}(u)\, du, \tag{S3}$$

and integrate $Y(t)$ over time, by applying Fubini's theorem:

$$\int_0^T Y(t)\, dt = \int_0^T \left(\int_0^t \hat{\psi}(u)\, du\right) dt = \int_0^T (T-u)\,\hat{\psi}(u)\, du, \tag{S4}$$

so that

$$\psi = \frac{2}{T^2}\int_0^T Y(t)\, dt. \tag{S5}$$

We define the treatment effect as the time-weighted difference between the temporal net growth rates. This corresponds to the scaled area between two curves (control and treatment):

$$\int_0^T \big(Y_{\text{ctrl}}(t) - Y_{\text{treat}}(t)\big)\, dt = \int_0^T (T-t)\,\big(\hat{\psi}_{\text{ctrl}}(t) - \hat{\psi}_{\text{treat}}(t)\big)\, dt, \tag{S6}$$

and is identical to the difference of the time-weighted net growth rates:

$$\tau = \psi_{\text{ctrl}} - \psi_{\text{treat}} = \frac{2}{T^2}\int_0^T \big(Y_{\text{ctrl}}(t) - Y_{\text{treat}}(t)\big)\, dt. \tag{S7}$$

**Intuition for $\psi$ and $\tau$.**

*Non-monophasic case.* $\tau$ represents a time-weighted average of instantaneous rate differences,

$$\tau = 2/T^2 \int_0^T (T-t)\,\big(\hat{\psi}_{\text{ctrl}}(t) - \hat{\psi}_{\text{treat}}(t)\big)\, dt,$$

where the weighting factor $(T-t)$ decreases linearly from $T$ to 0. Early rate differences thus contribute more strongly than later ones. This weighting is desirable because an early population reduction affects the population for longer than a delayed effect, even if both trajectories reach the same endpoint.

*Constant-rate case.* If the temporal net growth rate is constant, $\hat{\psi}(t) = \hat{\psi}$, then the time-weighted net growth rate (Equation S5) simplifies to $\psi = \hat{\psi}$.

**1B. Quantifying light noise.** We quantified stray light by using a calibration plate in which six source wells (E5, E12, E20, L5, L12, L20) contained 50 µL of luminescent overnight culture. All other wells were empty. A single luminescence read of all wells yielded raw intensities $\hat{I}$. We calculated the ratio of raw light intensities $\hat{I}_{\mathrm{e}}/\hat{I}_{\mathrm{l}}$ between empty wells $e$ and the nearest luminescent culture well $l$ as a function of the distance between their midpoints $d_{el}$ (with diagonal neighbours at a distance $\sqrt{2}$). For each discrete distance $d_{el}$, we summarised ratios by their median to obtain a kernel $r(d)$ (Figure S15). We estimate the light contribution from well $l$ into well $e$ as $\hat{I}_{el} = r(d_{el})\,\hat{I}_l$. For each experimental plate and time point, we corrected each well by subtracting received light noise,

$$I_e = \hat{I}_e - \sum_l r(d_{el})\,\hat{I}_l, \tag{S8}$$

where the sum is restricted to wells within $d \leq 3$.

#### 1C. Drug conditions and response functions.

**Single-drug pharmacodynamic curves.** For each drug we describe the relationship between concentration $c$ and net growth rate by the fitted pharmacodynamic curve

$$f(c) = \psi_0 - \frac{(\psi_0 - \psi_{\min})(c/\mathrm{zMIC})^{\kappa}}{(c/\mathrm{zMIC})^{\kappa} - (\psi_{\min}/\psi_0)}, \qquad [0,\infty) \to [\psi_{\min}, \psi_0]. \tag{S9}$$

**Hierarchical estimation.** Each drug's pharmacodynamic curve is estimated jointly across the experiments (one per drug pair) it appears in, using a Bayesian hierarchical model in a non-centred parameterisation. Each valid well of that drug's dilution series contributes its time-weighted net growth rate $\psi_w$ as one observation, modelled as $\psi_w \sim \mathcal{N}\left(f_{d,e}(c_w),\, \sigma_d^2\right)$, where $f_{d,e}$ is the Hill curve (Equation S9) of drug $d$ in experiment $e$, and $\sigma_d$ is a per-drug observation noise (Table S3). $\psi_0$ is pooled at the experiment level (shared by both drugs of a pair); $\kappa$ and zMIC are pooled per drug across that drug's five experiments. As $\psi_{\min}$ has to be smaller than the lowest rate observed for that drug ($\psi^*_{\min}$), we set $\psi_{\min} = 1.05\,\psi^*_{\min}$. $\psi_0$, $\kappa$ and zMIC are estimated by Markov chain Monte Carlo (NUTS [1]; PyMC [2]; four chains of 1000 draws after 1000 tuning steps; $\hat{R} \leq 1.01$, bulk effective sample size $> 1000$ [3]) and reported in Table S1.

**Closed-form inversion.** Using Equation S9, the inverse $f^{-1}$ can be written in closed form as

$$f^{-1}(\psi) = \mathrm{zMIC}\left(\frac{(\psi_0 - \psi)\,\psi_{\min}/\psi_0}{(\psi_0 - \psi) - (\psi_0 - \psi_{\min})}\right)^{1/\kappa}, \qquad \psi \in [\psi_{\min}, \psi_0]. \tag{S10}$$

**Cartesian conditions.** A *condition* is defined by a set of drug concentrations $\{c_i\}$, with drug $i \in \{\mathrm{AMO}, \mathrm{CHL}, \mathrm{COL}, \mathrm{PEN}, \mathrm{POL}, \mathrm{TET}\}$. The control condition corresponds to $\{c_i = 0\ \forall i\}$ and is denoted by $\emptyset$. For brevity, we write $X(c_A)$ when only one drug is present, $X(c_A, c_B)$ when two drugs are present, and $X(\emptyset)$ if no drug is present (control).

**Polar pharmacodynamic curves.** Conditions are reparameterised in polar coordinates $(z,\phi)$, where the combined dose $z = \sqrt{z_A^2 + z_B^2}$ and the mixing angle $\phi = \arctan 2(z_B, z_A)$ are computed from the zMIC-normalised concentrations $z_i = c_i/\mathrm{zMIC}_i$ (Methods). This yields the polar pharmacodynamic curve:

$$g(z,\phi) = \psi_0 - \frac{\left(\psi_0 - \psi_{\min}(\phi)\right)\left(z/\mathrm{zMIC}(\phi)\right)^{\kappa(\phi)}}{\left(z/\mathrm{zMIC}(\phi)\right)^{\kappa(\phi)} - \psi_{\min}(\phi)/\psi_0}. \tag{S11}$$

**1D. Bliss independence.** Bliss independence is defined by assuming that the probabilities of a cell being killed by drugs $A$ and $B$ within a fixed observation window are statistically independent [4]. Under this assumption,

$$p_{AB}^{\mathrm{Bliss}} = p_A + p_B - p_A\,p_B, \tag{S12}$$

where $p_A$ and $p_B$ denote the single-drug kill probabilities within $T$.

**Time–varying hazards.** Let $\hat{\tau}_i(t) \geq 0$ denote a time–dependent kill hazard under treatment $i$. The survival fraction at time $T$ is $S_i(T) = \exp\left(-\int_0^T \hat{\tau}_i(u)\,du\right)$. Bliss independence implies multiplicative survival $S_{AB}(T) = S_A(T)\,S_B(T)$, hence additive cumulative hazards:

$$\int_0^T \hat{\tau}_{AB}^{\mathrm{Bliss}}(u)\,du = \int_0^T \hat{\tau}_A(u)\,du + \int_0^T \hat{\tau}_B(u)\,du. \tag{S13}$$

To connect this survival formulation to the growth-based log-trajectories, we interpret the instantaneous treatment effect as the reduction in temporal net growth relative to the untreated control, i.e. $\hat{\tau}_i(t) = \hat{\psi}_{\mathrm{ctrl}}(t) - \hat{\psi}_i(t)$. Substituting $Y_i(t) = \int_0^t \hat{\psi}_i(u)\,du$ (Equation S3), it follows that:

$$Y_i(t) = Y_{\mathrm{ctrl}}(t) - \int_0^t \hat{\tau}_i(u)\,du.$$

Using additive cumulative hazards under Bliss then yields

$$Y_{AB}^{\text{Bliss}}(t) = Y_A(t) + Y_B(t) - Y_{\text{ctrl}}(t). \tag{S14}$$

Integrating Equation S14 over time yields the Bliss prediction for the treatment effect:

$$\tau^{\text{Bliss}}(c_A, c_B) = \tau(c_A) + \tau(c_B). \tag{S15}$$

**Bliss-based interaction score $\mu$.** Based on the Bliss expectation for a combination at concentrations $(c_A, c_B)$, we define the interaction score as the normalised divergence from the expectation:

$$\mu(c_A, c_B) = \frac{\tau(c_A) + \tau(c_B) - \tau(c_A, c_B)}{\tau(c_A, c_B)}. \tag{S16}$$

The score is only evaluated for conditions $(c_A, c_B)$ whose combined effect $\tau(c_A, c_B)$ is significantly greater than zero (5th percentile of its draws $> 0$).

> **Interpreting the Bliss interaction score $\mu$.**
> Negative values of $\mu$ indicate synergy ($\tau(c_A, c_B) > \tau(c_A) + \tau(c_B)$, i.e. the combination is more effective than the additive expectation), positive values indicate antagonism, and $\mu = 0$ corresponds to Bliss independence.

**1E. Loewe additivity.** Loewe additivity is a form of dose additivity, formalising self–additivity, i.e. that a drug combined with itself should behave like a higher dose of the same drug [5]. This concept is independent of the particular choice of effect measure, provided the effect can be represented on a shared, monotone dose–response scale. In our case, this effect scale is the time-weighted net growth rate $\psi$ (see Equation S5).

For a combination with the observed effect $\psi(c_A, c_B)$, Loewe additivity requires that the fractions of the equivalent single-drug concentrations sum to one,

$$\frac{c_A}{f_A^{-1}\big(\psi(c_A, c_B)\big)} + \frac{c_B}{f_B^{-1}\big(\psi(c_A, c_B)\big)} = 1, \tag{S17}$$

where $f_A$ and $f_B$ are the pharmacodynamic curves mapping concentrations to $\psi$, and $f_A^{-1}$ and $f_B^{-1}$ map an effect level $\psi$ to the corresponding equivalent single-drug concentration. All curve fits are shown in Figure S2. We later use the zMIC estimates for concentration normalisation, and the closed-form Hill inverses ($f_A^{-1}$, $f_B^{-1}$) to obtain the Loewe prediction $\psi_{\text{Loewe}}(c_A, c_B)$ by numerically solving Equation S17 for $\psi$.

**Loewe interaction score ($\nu$).** We quantify deviations from Loewe additivity using:

$$\nu(c_A, c_B) = \frac{c_A}{f_A^{-1}\big(\psi(c_A, c_B)\big)} + \frac{c_B}{f_B^{-1}\big(\psi(c_A, c_B)\big)} - 1. \tag{S18}$$

Because $f_A^{-1}$ and $f_B^{-1}$ are only defined on $\text{Im}(f_A) = [\psi_{\text{min},A}, \psi_0]$ and $\text{Im}(f_B) = [\psi_{\text{min},B}, \psi_0]$, $\nu$ is only defined if

$$\psi(c_A, c_B) \in \text{Im}(f_A) \cap \text{Im}(f_B) = [\max(\psi_{\text{min},A}, \psi_{\text{min},B}), \psi_0]. \tag{S19}$$

The score is only evaluated for conditions $(c_A, c_B)$ whose combined effect $\tau(c_A, c_B)$ is significantly greater than zero, as for $\mu$, and additionally only if $\nu$ is defined for at least 10% of posterior draws.

> **Interpreting the Loewe interaction score $\nu$.**
> By construction $\nu = 0$ corresponds to Loewe additivity, $\nu < 0$ indicates synergy, and $\nu > 0$ indicates antagonism. Because Loewe additivity requires equivalent single-drug concentrations for both drugs at the observed effect level, $\nu$ can only be evaluated when $\psi(c_A, c_B)$ lies in the overlap of the two single-drug effect ranges, i.e. $\psi \geq \max(\psi_{\text{min},A}, \psi_{\text{min},B})$.

**1F. Peptide–antibiotic interaction model.** We consider two drugs, $A$ and $B$, acting on a population with a size trajectory $x(t)$. Drug $A$ represents a peptide-like effect that causes an instantaneous multiplicative reduction in population size by a factor $\alpha \in (0, 1]$, whereas drug $B$ represents a conventional antibiotic that changes the subsequent net growth rate but does not cause an initial drop. We assume that the combination of two drugs induces a sharp drop by the factor $\alpha$ followed by net growth, determined by drug B:

$$\begin{aligned} x_{\text{ctrl}}(t) &= x_0 \exp\big(\hat{\psi}_{\text{ctrl}}\, t\big), \\ x_B(t) &= x_0 \exp\big(\hat{\psi}_B\, t\big), \\ x_A(t) &= \alpha\, x_{\text{ctrl}}(t), \\ x_{AB}(t) &= \alpha\, x_B(t), \end{aligned}$$

and:

$$\begin{aligned} Y_{\text{ctrl}}(t) &= \ln x_0 + \hat{\psi}_{\text{ctrl}}\, t, \\ Y_B(t) &= \ln x_0 + \hat{\psi}_B\, t, \\ Y_A(t) &= Y_{\text{ctrl}}(t) + \ln\alpha, \\ Y_{AB}(t) &= Y_B(t) + \ln\alpha. \end{aligned}$$

Using the definition of the treatment effect $\tau_i = \frac{2}{T^2}\int_0^T \big(Y_{\text{ctrl}}(t) - Y_i(t)\big)\,dt$, we obtain the following.

$$\tau_A = \frac{2}{T^2}\int_0^T (-\ln\alpha)\,dt, \qquad \tau_B = \frac{2}{T^2}\int_0^T \big(\hat{\psi}_{\text{ctrl}} - \hat{\psi}_B\big)t\,dt,$$

and

$$\tau_{AB} = \frac{2}{T^2}\int_0^T \Big(\big(\hat{\psi}_{\text{ctrl}} - \hat{\psi}_B\big)t - \ln\alpha\Big)\,dt. \tag{S20}$$

Separating the two terms in Equation S20 yields

$$\tau_{AB} = \tau_A + \tau_B, \tag{S21}$$

which matches the Bliss expectation for the combined treatment effect.

#### 1G. Interaction inference on the checkerboard conditions.

**Interaction field.** We measure the treatment effect of each checkerboard condition with four replicates. For conditions in which the population declines rapidly, even fewer replicates remain eligible, as replicates falling below the detection limit are excluded. Estimated on their own, such conditions are not only poorly informed but can also appear more certain than they are (a single replicate has no scatter at all). Additionally, the surviving replicates are a selective sample, as the fastest-declining ones are the most likely to be excluded, which biases the estimate towards weaker killing. At the same time, neighbouring conditions differ by a single dilution step, and their effects are expected to be similar. We therefore estimate all conditions of a checkerboard together, as one Gaussian Markov random field [6], so that every condition is informed primarily by its own replicates, but also by its immediate neighbours. We fit the field using Gibbs sampling, a standard method for sampling a multivariate distribution. Each condition is drawn in turn from its conditional distribution, given its own replicates and the current values of its neighbours. One pass over all conditions is a sweep, and the updated values after a sweep are one posterior draw of the field. Indexing the conditions of one checkerboard by $i$, we write $\psi_i$ for the field's net growth rate at condition $i$ and $y_{ik}$ for the estimate from its $k$-th valid replicate. Replicates are modelled as $y_{ik} \sim \mathcal{N}(\psi_i, \sigma_i^2)$, and conditions adjacent on the dose grid are tied by $\psi_i - \psi_j \sim \mathcal{N}(0, \omega_\psi^2)$, where $\omega_\psi$ sets how strongly neighbours are smoothed (Table S3). The per-condition scale $\sigma_i$ is computed once before sampling, from the within-condition scatter smoothed across neighbours on the log scale (Table S3), and held fixed throughout the Gibbs run. Below-detection-limit replicates and conditions with no valid replicate are excluded, along with pockets of conditions that become disconnected from the main field as a result.

**Regime-level random-effects meta-analysis.** To summarise interactions for a given drug pair within a regime (sub-inhibitory or inhibitory), we aggregate the eligible conditions' score distributions via a Bayesian random-effects meta-analysis [7]. We evaluate its posterior in a Python implementation on a dense grid (600 points for $M$, 200 for $\sigma$, spanning the range of the contributing conditions). Because the model has only two unknowns, $M$ and $\sigma$, evaluating the posterior on a grid is much faster than sampling it. A condition contributes to a regime only if at least 5% of the posterior draws place it in that regime and yield a defined score. Each eligible condition $i$ contributes its posterior median score $y_i$ (i.e. $\mu_i$ or $\nu_i$) and the standard deviation $s_i$ of that score. Because $\pm 1$ standard deviation around the mean of a Gaussian spans $\approx 68\%$ of its mass, we obtain $s_i$ as half the distance between the 16th and 84th percentiles of the draws. These are modelled as

$$y_i \sim \mathcal{N}(M,\, s_i^2/w_i + \sigma^2), \qquad M \sim \mathcal{N}(0,1), \qquad \sigma \sim \text{HalfNormal}(1), \tag{S22}$$

where $M$ is the regime-level score, $\sigma$ the between-condition heterogeneity, and $w_i$ each condition's mixing weight ($w_i = 1$ in the reported, unweighted analysis; $w_i = \sin(2\phi_i)$ is a sensitivity variant). For the between-condition standard deviation $\sigma$ we use a weak, fixed HalfNormal$(1)$, since no raw data estimates it ahead of the fit. We report the posterior median and central 90% interval of $M$ (main-text Fig 3, Table S2). We validated the Python implementation against `bayesmeta` [7], the R package implementing the same model, by running both on the same per-condition inputs. The posterior medians and interval bounds differed by at most 0.017 (median $5 \times 10^{-5}$), and no interaction label changed. The comparison is part of the code deposit (DOI: 10.5281/zenodo.18374151).

**Regime alignment.** We assess whether the posterior distributions of the inhibitory and sub-inhibitory regimes align by inferring the distribution of their difference, $D = M_{\text{inh}} - M_{\text{sub}}$, and asking how much of it lies on one side of zero. We report not significant (n.s.) below 95%, significant ($*$) at 95%, and significant ($**$) at 99%.

**1H. Robustness of the interaction labels.** Next, we assess how sensitive the regime-level interaction labels are to our analysis choices. The base analysis used throughout the manuscript uses the time-weighted net growth rate $\psi$ (Equation S5) over the window $T \approx 2\,\mathrm{h}$, aggregated via the random-effects meta-analysis of subsection 1G. Here, we repeat the regime-wise interaction analysis, changing one of these choices at a time. For each configuration we recomputed the interaction scores and their regime-level median and central 90% interval exactly as in subsection 1G, for both reference models (Bliss $\mu$ and Loewe $\nu$) and both regimes (sub-inhibitory and inhibitory).

**Time-window sensitivity.** We re-estimated all rates over the shorter window $T = 1$, leaving the aggregation method otherwise unchanged. Because validity is assessed over $[0, T]$, the shorter window additionally retains wells that fall below the detection limit only after $t = 1$. In Figure S8 we plot the base-window score ($T = 2$, $x$) against the shorter-window score ($T = 1$, $y$). Proximity to the $y = x$ diagonal indicates that the regime posterior is time-invariant. Shortening the window changed 17 of the 60 regime labels (15 pairs $\times$ 2 regimes $\times$ 2 reference models), and inverted one: COL+PEN under Bliss in the inhibitory regime, from antagonistic to synergistic.

**Methodological sensitivity.** We tested five further analysis choices, each changing one element of the base analysis:

**Weighting (weighted).** We repeated the base aggregation with each condition weighted by $\sin(2\phi)$, emphasising conditions with near-equal effect ratios and down-weighting conditions near the pure-drug edges.

**Aggregation method (pool).** We pooled every eligible condition's score draws into one combined sample and read the median and interval directly off its quantiles, in place of the random-effects meta-analysis; the resulting interval reflects the spread across conditions.

**Aggregation method (boot).** For each posterior draw we took the median score across the regime's conditions, resampling those conditions with replacement (a bootstrap) so that variability between conditions enters the estimate. This gives one value per draw, and we read the median and interval off their spread, in place of the random-effects meta-analysis.

**Rate estimator (slope).** Instead of the time-weighted $\psi$, we estimated each per-well net growth rate as the ordinary-least-squares slope of the log-normalised signal $Y(t) = \ln(I(t)/I(0))$.

**Condition sampling (sector 1–3).** We re-aggregated the base estimation but stratified the conditions into three equally sized $24^\circ$ mixing-angle sectors spanning 9°–81° ($\phi \in [9^\circ, 33^\circ]$, $[33^\circ, 57^\circ]$, $[57^\circ, 81^\circ]$), sampling uniformly within each sector and summarising each separately.

We summarise the results in one grid per reference model, Bliss (Figure S9) and Loewe (Figure S10).

#### 1I. Priors.

**Prior calibration.** Every prior width in the hierarchical model is a multiplier times an empirical base estimate of that quantity's natural between-experiment (or residual) variability, computed once from the raw mono dose–response data, independently of the fitted model (Table S3). The multiplier is $5\times$ for population-mean (location) parameters and $\sigma$. We use a tighter $2\times$ for the between-experiment spread hyperparameters ($s_{\psi_0}$, $s_{\log\kappa,d}$, $s_{\log \mathrm{zMIC},d}$), whose base estimate directly represents the quantity itself, to preserve the partial-pooling shrinkage.

**Prior sensitivity.** To check how the chosen priors affect their posteriors, we repeated the analysis twice, once with all priors widened and once with all priors tightened by a factor of two (Table S4). The pharmacodynamic and regime-pooling posteriors are mostly unaffected, moving by at most 0.05 standard deviations at the median. The field's smoothing priors move conditions with four valid replicates by 0.15, and those with fewer by about one standard deviation. This shift is expected and intentional, as the smoothing scales $\omega_\psi$ and $\omega_{\log\sigma}$ regulate how strongly conditions share information, on which weakly informed conditions depend heavily.

#### 1J. Surface analyses.

**Continuous surface construction.** For each pair of drugs $(A, B)$, we use 100 draws of the interaction field for that pair's checkerboard (subsection 1G). We draw-wise apply an isotonic regression to every row and column of the grid, making $\psi$ strictly decreasing in each drug's concentration (`sklearn.isotonic.IsotonicRegression` [8]). Monotonicity ensures unique invertibility, necessary to calculate the predictions for Loewe and to estimate the isoboles. On this monotone grid we fit a continuous surface using a cubic (PCHIP) interpolation (`scipy.interpolate.PchipInterpolator` [9]). For each drug combination we calculate a consensus surface $\psi^{\mathrm{cons}}$ as the pointwise median across these per-draw surfaces, and all quantities derived from it (Bliss and Loewe predictions, isoboles, polar pharmacodynamic curves, angular-dependence curves), each computed per draw and then aggregated by median.

**Surface domain.** We construct the concentration grid only from single-drug conditions with at least one valid replicate. The interpolation is then confined to regions spanned by consecutively valid conditions. For the quantities derived from the surface (Bliss and Loewe predictions, isoboles, polar pharmacodynamic curves, angular-dependence curves) we additionally discard points at which fewer than half of the draws yield a finite value.

**Uncertainty estimation.** We construct 100 surfaces: for each, we draw one $\psi$ per condition and add Gaussian noise with that condition's previously estimated standard deviation $\sigma_i$ (subsection 1G), then fit a surface through the resulting values as above. Uncertainty bands in Figure S12 and Figure S14 are percentiles across these surfaces.

**1K. Single-drug inoculum effect analysis.** During data analysis, we noted that inocula varied more than expected, which we traced to variability induced by the pintool. Across all wells, the pre-treatment luminescence had a mean of $\langle I_0 \rangle = 4.61 \times 10^4$ RLU and a standard deviation of $\sigma(I_0) = 6.89 \times 10^4$ RLU, which decomposes into a between-experiment component $\sigma_{\text{between}} = 4.33 \times 10^4$ RLU and a within-experiment component $\sigma_{\text{within}} = 5.47 \times 10^4$ RLU. To investigate whether the observed net growth rates $\psi$ systematically depended on inoculum size, we used the pre-treatment luminescence signal ($I_0$) as a proxy for initial cell density.

For all single-drug conditions, we fitted an ordinary least-squares regression of $\psi$ on $I_0$ to obtain a concentration-specific slope $\gamma_{\text{inoculum}}(c_i)$ together with a confidence interval, the $p$-value and the coefficient of determination $R^2$. These slopes and their uncertainty summaries were visualised as concentration–slope profiles with 95% intervals in Figure S16. Across AMO, CHL, PEN, and TET, the estimated slopes were negligible across concentrations, whereas COL and POL showed the strongest dependence on $I_0$ at intermediate concentrations (around 1 $\frac{\mu g}{mL}$).

To assess whether inoculum size varies systematically along the one-dimensional concentration index, we pooled all single-drug wells and mapped each well to its concentration index. We then performed a permutation test by permuting the light intensity across the concentration indices to obtain a null distribution. We found no evidence for a systematic trend of inoculum size with index in our data ($p = 0.092$), suggesting that inoculum variation contributes noise but does not introduce a strong directional bias along the concentration series. It therefore contributes mainly to the increased scatter in the COL and POL single-drug pharmacodynamic curves (Figure S2).

183 ## SI tables

**Table S1:** Hierarchical pharmacodynamic model parameters (Equation S9) for each drug: posterior median and 95% credible interval of $\kappa$ and zMIC, together with their between-experiment coefficient of variation (CV). $\psi_0$ (the drug-free rate) is shared by both drugs of an experiment. Its global population estimate is $1.84$ [1.81, 1.87] (95% CI), and between experiments it varies with a standard deviation of $0.043$ (CV $2.3\%$). For $\psi_{\min}$ we only know that it lies at or below the lowest rate observed for that drug, $\psi^*_{\min}$; accordingly we set $\psi_{\min} = 1.05\,\psi^*_{\min}$, just below that value.

| drug | $\psi_{\min}$ | $\kappa$ | zMIC | $CV_\kappa$ | $CV_z$ |
|---|---|---|---|---|---|
| AMO | -4.55 | 1.35 [1.12, 1.62] | 8.28 [7.89, 8.64] | 15% | 2% |
| CHL | -1.25 | 0.78 [0.66, 0.92] | 60.39 [48.08, 75.44] | 15% | 20% |
| COL | -8.41 | 1.48 [1.10, 2.08] | 0.60 [0.43, 0.85] | 25% | 31% |
| PEN | -4.76 | 1.01 [0.82, 1.25] | 153.44 [138.16, 171.75] | 17% | 9% |
| POL | -8.07 | 1.06 [0.69, 1.74] | 0.49 [0.35, 0.69] | 33% | 29% |
| TET | -0.43 | 0.92 [0.87, 0.98] | 15.31 [12.96, 17.97] | 2% | 14% |

**Table S2:** Interaction labels (N: no interaction (independent), S: synergistic, A: antagonistic) for each drug combination based on Bliss independence ($\mu$) and Loewe additivity ($\nu$), evaluated separately for sub-inhibitory and inhibitory concentration regimes. A regime is labelled synergistic or antagonistic when at least $95\%$ of its regime-level posterior lies on that side of zero, and independent otherwise (Bayesian random-effects meta-analysis, subsection 1G).

| Drug A | Drug B | Bliss | | Loewe | |
|---|---|---|---|---|---|
| | | sub-inhibitory | inhibitory | sub-inhibitory | inhibitory |
| AMO | CHL | A | A | A | N |
| AMO | COL | S | A | A | A |
| AMO | PEN | S | A | N | A |
| AMO | POL | N | A | A | A |
| AMO | TET | A | A | A | S |
| CHL | COL | S | S | N | S |
| CHL | PEN | A | A | A | A |
| CHL | POL | S | S | S | S |
| CHL | TET | S | A | S | S |
| COL | PEN | N | A | A | A |
| COL | POL | N | A | A | A |
| COL | TET | N | S | A | N |
| PEN | POL | N | A | A | N |
| PEN | TET | A | A | A | N |
| POL | TET | N | S | A | N |

**Table S3:** All priors used in the analysis. The pharmacodynamic priors are centred and scaled from the raw dose–response data (subsection 1I); the field and meta-analysis priors are fixed. *Centres* are preliminary estimates of the parameter itself, marked by a superscript $(0)$: $\psi_0^{(0)} = 1.695$ is the mean $\psi$ across the $60$ control wells, and $\kappa_d^{(0)}$, $\mathrm{zMIC}_d^{(0)}$ come from a per-drug least-squares Hill fit. *Scales* are the product of a base estimate $b$ and a multiplier, $5\times$ for location parameters and observation noise and $2\times$ for between-experiment spreads. Base estimates: $b_{\psi_0} = 0.047$ (between-experiment SD of control $\psi$, 15 pairs), $b_\kappa = 0.327$ and $b_{\mathrm{zMIC}} = 0.182$ (pooled between-experiment SD of $\log\kappa$ and $\log \mathrm{zMIC}$, 6 drugs), $b_\sigma = 0.400$ (pooled residual SD of per-drug least-squares Hill fits).

| Prior | Centre | Scale | Comment |
|---|---|---|---|
| **Hierarchical pharmacodynamic model (Equation S9)** | | | |
| $\beta_{\psi_0} \sim \mathcal{N}\left(\psi_0^{(0)},\ (5b_{\psi_0})^2\right)$ | 1.695 | 0.235 | Population mean of the drug-free rate, informed by 60 control wells |
| $\beta_{\log\kappa,d} \sim \mathcal{N}\left(\log\kappa_d^{(0)},\ (5b_\kappa)^2\right)$ | per drug | 1.635 | Population mean of $\log\kappa$, informed by five experiments per drug |
| $\beta_{\log \mathrm{zMIC},d} \sim \mathcal{N}\left(\log \mathrm{zMIC}_d^{(0)},\ (5b_{\mathrm{zMIC}})^2\right)$ | per drug | 0.91 | Population mean of $\log \mathrm{zMIC}$, informed by five experiments per drug |
| $s_{\psi_0} \sim \mathrm{HalfNormal}(2b_{\psi_0})$ | — | 0.094 | Between-experiment spread of $\psi_0$, shared by both drugs of an experiment |
| $s_{\log\kappa,d} \sim \mathrm{HalfNormal}(2b_\kappa)$ | — | 0.654 | Between-experiment spread of $\log\kappa$, per drug |
| $s_{\log \mathrm{zMIC},d} \sim \mathrm{HalfNormal}(2b_{\mathrm{zMIC}})$ | — | 0.364 | Between-experiment spread of $\log \mathrm{zMIC}$, per drug |
| $\sigma_d \sim \mathrm{HalfNormal}(5b_\sigma)$ | — | 2.0 | Residual observation noise per drug; scaled $5\times$ because it is not a hierarchical spread |
| **Checkerboard $\psi$ field (subsection 1G)** | | | |
| $\psi_i - \psi_j \sim \mathcal{N}\left(0,\ \omega_\psi^2\right)$ | 0 | 0.5 | Difference between conditions adjacent on the dose grid (four-neighbourhood); the only prior acting on the interior, as the single-drug edges are fixed |
| $\log\sigma_i - \log\sigma_j \sim \mathcal{N}\left(0,\ \omega_{\log\sigma}^2\right)$ | 0 | 1.0 | Difference between adjacent conditions of the log noise scale, smoothing the per-condition noise scale so that conditions with few valid replicates do not receive extreme weights |
| **Regime-aggregation meta-analysis (Equation S22)** | | | |
| $M \sim \mathcal{N}(0,\ 1)$ | 0 | 1 | Regime-level pooled score; symmetric about 0, so synergy and antagonism are equally likely a priori |
| $\sigma_M \sim \mathrm{HalfNormal}(1)$ | — | 1 | Between-condition spread; not calibrated from a base estimate, as none exists before the fit |

**Table S4:** Prior sensitivity of the three models. Every prior was widened and tightened by a factor of two. Each entry summarises how far posterior medians move under that change, in units of their own posterior standard deviation: the median and the 95th percentile across the quantities in that row. The pharmacodynamic and regime-pooling priors have little influence on their posteriors (median shifts $< 0.05$). The field's priors smooth across neighbouring conditions, so we report its conditions separately by their number of valid replicates: those with four move by $\approx 0.15$ and those with fewer by $\approx 1$ standard deviation.

| Model | Prior variation | Valid replicates | Quantities | Median shift | 95th percentile |
|---|---|---|---:|---|---|
| Pharmacodynamic curves | all priors $\times 0.5$ | all | 90 | 0.050 | 0.169 |
| Pharmacodynamic curves | all priors $\times 2$ | all | 90 | 0.031 | 0.084 |
| Interaction field | all priors $\times 0.5$ | 4 | 1358 | 0.150 | 1.436 |
| Interaction field | all priors $\times 0.5$ | 1–3 | 112 | 1.204 | 3.805 |
| Interaction field | all priors $\times 2$ | 4 | 1358 | 0.040 | 0.974 |
| Interaction field | all priors $\times 2$ | 1–3 | 112 | 1.111 | 3.689 |
| Regime pooling | all priors $\times 0.5$ | all | 60 | 0.036 | 0.131 |
| Regime pooling | all priors $\times 2$ | all | 60 | 0.009 | 0.041 |

**Table S5:** DMSO content (%) of every checkerboard condition of CHL+TET, for which both components are dissolved in DMSO. Rows and columns are the concentration indices, from drug-free (0) to the highest concentration (11). Cells with $\geq 1.25\%$ DMSO are shown in red.

| DMSO (columns) / DMSO (rows) | 0 | 1 | 2 | 3 | 4 | 5 | 6 | 7 | 8 | 9 | 10 | 11 |
|---|---|---|---|---|---|---|---|---|---|---|---|---|
| 0 | 0 | $\frac{5}{1024}$ | $\frac{5}{512}$ | $\frac{5}{256}$ | $\frac{5}{128}$ | $\frac{5}{64}$ | $\frac{5}{32}$ | $\frac{5}{16}$ | $\frac{5}{8}$ | $\frac{5}{4}$ | $\frac{5}{2}$ | 5 |
| 1 | $\frac{5}{1024}$ | $\frac{5}{512}$ | $\frac{15}{1024}$ | $\frac{25}{1024}$ | $\frac{45}{1024}$ | $\frac{85}{1024}$ | $\frac{165}{1024}$ | $\frac{325}{1024}$ | $\frac{645}{1024}$ | $\frac{1285}{1024}$ | $\frac{2565}{1024}$ | $\frac{5125}{1024}$ |
| 2 | $\frac{5}{512}$ | $\frac{15}{1024}$ | $\frac{5}{256}$ | $\frac{15}{512}$ | $\frac{25}{512}$ | $\frac{45}{512}$ | $\frac{85}{512}$ | $\frac{165}{512}$ | $\frac{325}{512}$ | $\frac{645}{512}$ | $\frac{1285}{512}$ | $\frac{2565}{512}$ |
| 3 | $\frac{5}{256}$ | $\frac{25}{1024}$ | $\frac{15}{512}$ | $\frac{5}{128}$ | $\frac{15}{256}$ | $\frac{25}{256}$ | $\frac{45}{256}$ | $\frac{85}{256}$ | $\frac{165}{256}$ | $\frac{325}{256}$ | $\frac{645}{256}$ | $\frac{1285}{256}$ |
| 4 | $\frac{5}{128}$ | $\frac{45}{1024}$ | $\frac{25}{512}$ | $\frac{15}{256}$ | $\frac{5}{64}$ | $\frac{15}{128}$ | $\frac{25}{128}$ | $\frac{45}{128}$ | $\frac{85}{128}$ | $\frac{165}{128}$ | $\frac{325}{128}$ | $\frac{645}{128}$ |
| 5 | $\frac{5}{64}$ | $\frac{85}{1024}$ | $\frac{45}{512}$ | $\frac{25}{256}$ | $\frac{15}{128}$ | $\frac{5}{32}$ | $\frac{15}{64}$ | $\frac{25}{64}$ | $\frac{45}{64}$ | $\frac{85}{64}$ | $\frac{165}{64}$ | $\frac{325}{64}$ |
| 6 | $\frac{5}{32}$ | $\frac{165}{1024}$ | $\frac{85}{512}$ | $\frac{45}{256}$ | $\frac{25}{128}$ | $\frac{15}{64}$ | $\frac{5}{16}$ | $\frac{15}{32}$ | $\frac{25}{32}$ | $\frac{45}{32}$ | $\frac{85}{32}$ | $\frac{165}{32}$ |
| 7 | $\frac{5}{16}$ | $\frac{325}{1024}$ | $\frac{165}{512}$ | $\frac{85}{256}$ | $\frac{45}{128}$ | $\frac{25}{64}$ | $\frac{15}{32}$ | $\frac{5}{8}$ | $\frac{15}{16}$ | $\frac{25}{16}$ | $\frac{45}{16}$ | $\frac{85}{16}$ |
| 8 | $\frac{5}{8}$ | $\frac{645}{1024}$ | $\frac{325}{512}$ | $\frac{165}{256}$ | $\frac{85}{128}$ | $\frac{45}{64}$ | $\frac{25}{32}$ | $\frac{15}{16}$ | $\frac{5}{4}$ | $\frac{15}{8}$ | $\frac{25}{8}$ | $\frac{45}{8}$ |
| 9 | $\frac{5}{4}$ | $\frac{1285}{1024}$ | $\frac{645}{512}$ | $\frac{325}{256}$ | $\frac{165}{128}$ | $\frac{85}{64}$ | $\frac{45}{32}$ | $\frac{25}{16}$ | $\frac{15}{8}$ | $\frac{5}{2}$ | $\frac{15}{4}$ | $\frac{25}{4}$ |
| 10 | $\frac{5}{2}$ | $\frac{2565}{1024}$ | $\frac{1285}{512}$ | $\frac{645}{256}$ | $\frac{325}{128}$ | $\frac{165}{64}$ | $\frac{85}{32}$ | $\frac{45}{16}$ | $\frac{25}{8}$ | $\frac{15}{4}$ | 5 | $\frac{15}{2}$ |
| 11 | 5 | $\frac{5125}{1024}$ | $\frac{2565}{512}$ | $\frac{1285}{256}$ | $\frac{645}{128}$ | $\frac{325}{64}$ | $\frac{165}{32}$ | $\frac{85}{16}$ | $\frac{45}{8}$ | $\frac{25}{4}$ | $\frac{15}{2}$ | 10 |

**Table S6:** DMSO content (%) for the pairs in which only one component is dissolved in DMSO, the other in water. Rows and columns are the concentration indices, from drug-free (0) to the highest concentration (11). Cells with $\geq 1.25\%$ DMSO are shown in red.

| DMSO (columns) / $H_2O$ (rows) | 0 | 1 | 2 | 3 | 4 | 5 | 6 | 7 | 8 | 9 | 10 | 11 |
|---|---|---|---|---|---|---|---|---|---|---|---|---|
| 0 | 0 | $\frac{5}{1024}$ | $\frac{5}{512}$ | $\frac{5}{256}$ | $\frac{5}{128}$ | $\frac{5}{64}$ | $\frac{5}{32}$ | $\frac{5}{16}$ | $\frac{5}{8}$ | $\frac{5}{4}$ | $\frac{5}{2}$ | 5 |
| 1 | 0 | $\frac{5}{1024}$ | $\frac{5}{512}$ | $\frac{5}{256}$ | $\frac{5}{128}$ | $\frac{5}{64}$ | $\frac{5}{32}$ | $\frac{5}{16}$ | $\frac{5}{8}$ | $\frac{5}{4}$ | $\frac{5}{2}$ | 5 |
| 2 | 0 | $\frac{5}{1024}$ | $\frac{5}{512}$ | $\frac{5}{256}$ | $\frac{5}{128}$ | $\frac{5}{64}$ | $\frac{5}{32}$ | $\frac{5}{16}$ | $\frac{5}{8}$ | $\frac{5}{4}$ | $\frac{5}{2}$ | 5 |
| 3 | 0 | $\frac{5}{1024}$ | $\frac{5}{512}$ | $\frac{5}{256}$ | $\frac{5}{128}$ | $\frac{5}{64}$ | $\frac{5}{32}$ | $\frac{5}{16}$ | $\frac{5}{8}$ | $\frac{5}{4}$ | $\frac{5}{2}$ | 5 |
| 4 | 0 | $\frac{5}{1024}$ | $\frac{5}{512}$ | $\frac{5}{256}$ | $\frac{5}{128}$ | $\frac{5}{64}$ | $\frac{5}{32}$ | $\frac{5}{16}$ | $\frac{5}{8}$ | $\frac{5}{4}$ | $\frac{5}{2}$ | 5 |
| 5 | 0 | $\frac{5}{1024}$ | $\frac{5}{512}$ | $\frac{5}{256}$ | $\frac{5}{128}$ | $\frac{5}{64}$ | $\frac{5}{32}$ | $\frac{5}{16}$ | $\frac{5}{8}$ | $\frac{5}{4}$ | $\frac{5}{2}$ | 5 |
| 6 | 0 | $\frac{5}{1024}$ | $\frac{5}{512}$ | $\frac{5}{256}$ | $\frac{5}{128}$ | $\frac{5}{64}$ | $\frac{5}{32}$ | $\frac{5}{16}$ | $\frac{5}{8}$ | $\frac{5}{4}$ | $\frac{5}{2}$ | 5 |
| 7 | 0 | $\frac{5}{1024}$ | $\frac{5}{512}$ | $\frac{5}{256}$ | $\frac{5}{128}$ | $\frac{5}{64}$ | $\frac{5}{32}$ | $\frac{5}{16}$ | $\frac{5}{8}$ | $\frac{5}{4}$ | $\frac{5}{2}$ | 5 |
| 8 | 0 | $\frac{5}{1024}$ | $\frac{5}{512}$ | $\frac{5}{256}$ | $\frac{5}{128}$ | $\frac{5}{64}$ | $\frac{5}{32}$ | $\frac{5}{16}$ | $\frac{5}{8}$ | $\frac{5}{4}$ | $\frac{5}{2}$ | 5 |
| 9 | 0 | $\frac{5}{1024}$ | $\frac{5}{512}$ | $\frac{5}{256}$ | $\frac{5}{128}$ | $\frac{5}{64}$ | $\frac{5}{32}$ | $\frac{5}{16}$ | $\frac{5}{8}$ | $\frac{5}{4}$ | $\frac{5}{2}$ | 5 |
| 10 | 0 | $\frac{5}{1024}$ | $\frac{5}{512}$ | $\frac{5}{256}$ | $\frac{5}{128}$ | $\frac{5}{64}$ | $\frac{5}{32}$ | $\frac{5}{16}$ | $\frac{5}{8}$ | $\frac{5}{4}$ | $\frac{5}{2}$ | 5 |
| 11 | 0 | $\frac{5}{1024}$ | $\frac{5}{512}$ | $\frac{5}{256}$ | $\frac{5}{128}$ | $\frac{5}{64}$ | $\frac{5}{32}$ | $\frac{5}{16}$ | $\frac{5}{8}$ | $\frac{5}{4}$ | $\frac{5}{2}$ | 5 |

### SI figures

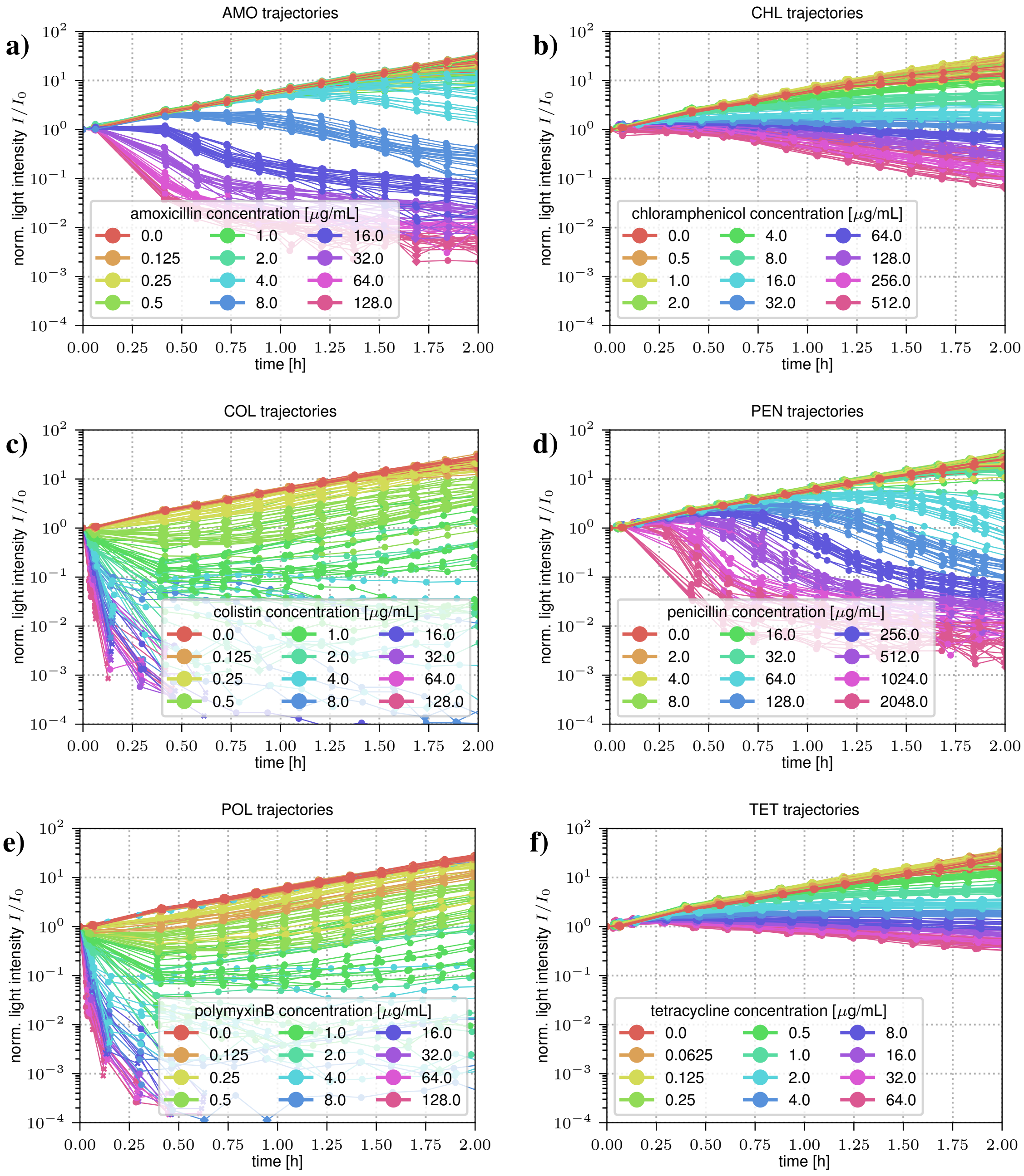


**Figure S1:** Normalised luminescence trajectories $I(t)/I(0)$ for single-drug treatments. Panels (a–f) show trajectories across $12$ single-drug concentrations for amoxicillin (AMO), chloramphenicol (CHL), colistin (COL), penicillin (PEN), polymyxin B (POL), and tetracycline (TET), respectively. For each concentration, we show all $20$ replicate curves obtained from five independent combination experiments (per drug) with four biological replicates each. For trajectories that fall temporarily (less than 20% of datapoints) below the detection limit, we substituted the respective timepoints with the detection limit (diamonds). The last regular datapoint of an invalid trajectory (more than 20% below the detection limit) is indicated with an "x", and all other datapoints are shown as circles.

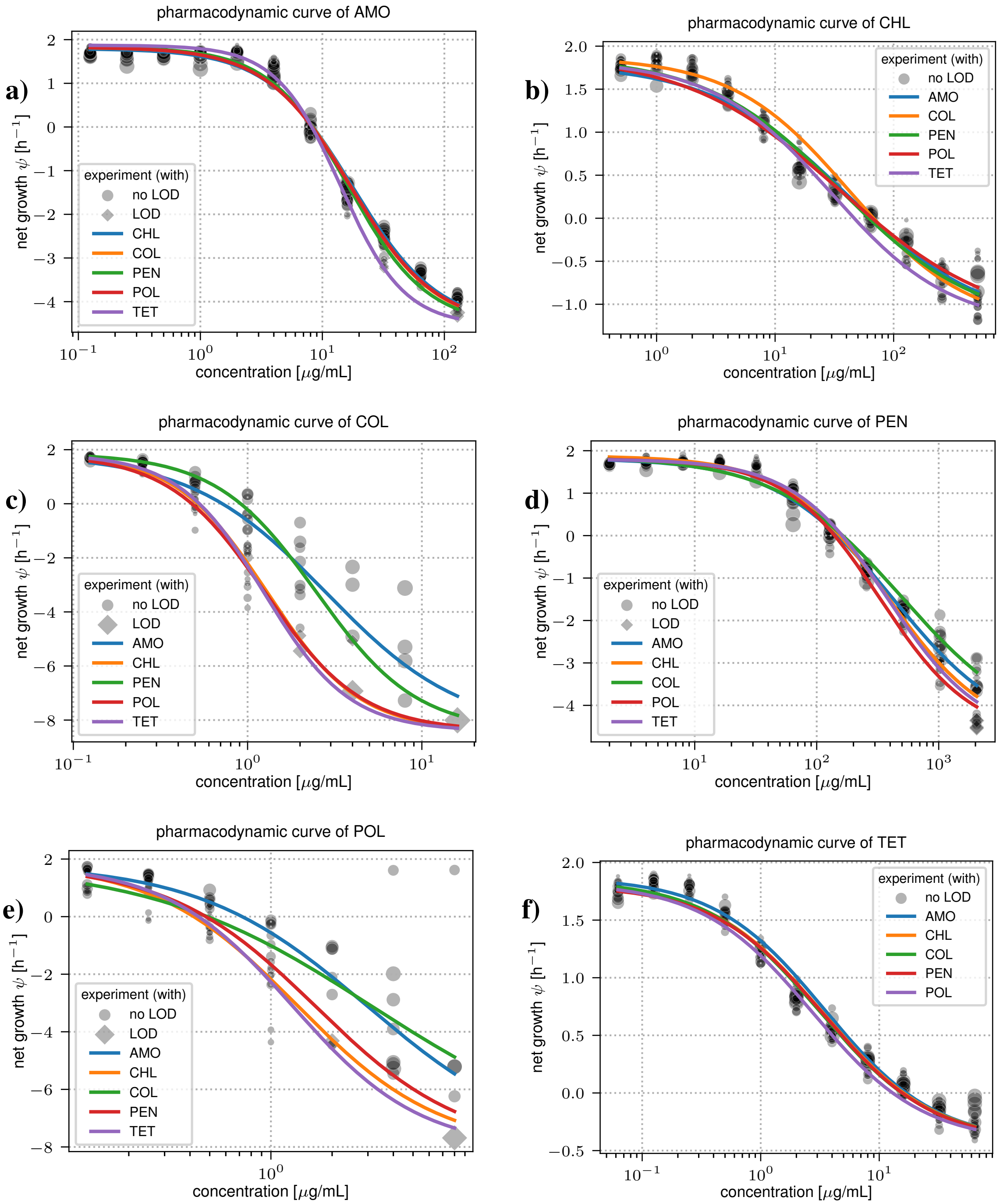


**Figure S2:** Panels (a–f) show the single-drug pharmacodynamic curves corresponding to the timecourse data for amoxicillin (AMO), chloramphenicol (CHL), colistin (COL), penicillin (PEN), polymyxin B (POL), and tetracycline (TET), respectively. For each concentration, we aggregate time-weighted net growth rate estimates from 20 replicates obtained from five independent combination experiments (per drug) with four biological replicates each. We included only "valid" estimates, defined as trajectories with less than 20% of timepoints below the LoD. Valid estimates containing LoD-censored timepoints are shown as diamonds, and estimates without LoD-censored timepoints are shown as circles. Marker size is scaled with $\log_{10}$ of the inoculum $I(0)$. Curves show the per-experiment posterior-mean pharmacodynamic fits from the hierarchical Bayesian model (subsection 1C).

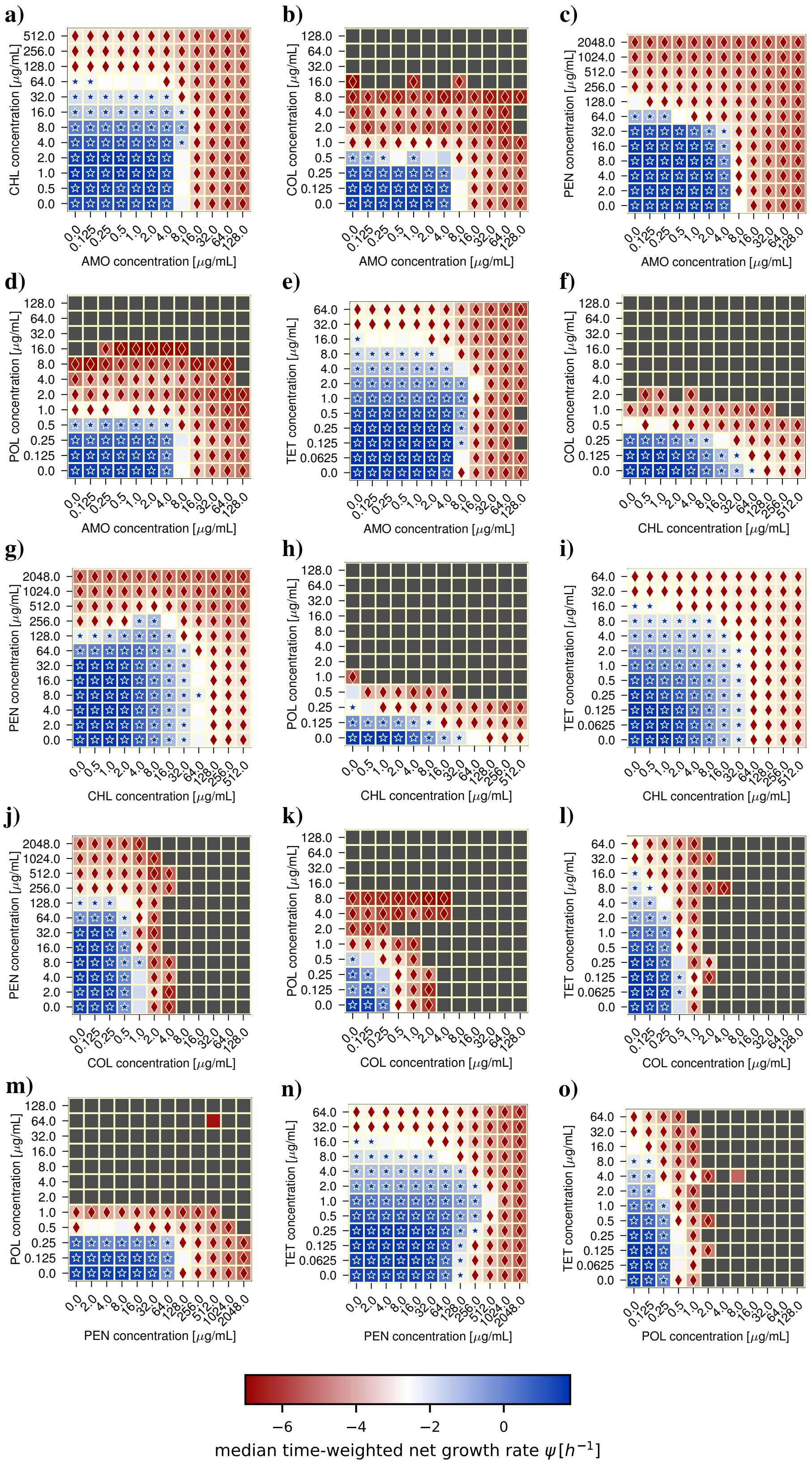


**Figure S3:** Median time-weighted net growth rate $\psi$ across all drug combinations measured in checkerboard assays. Each panel corresponds to one drug pair, and each cell represents a condition $(c_A, c_B)$. Colours indicate the median $\psi$ across biological replicates. Stars denote conditions that are predominantly ($\geq 95\%$ of the posterior) sub-inhibitory (net growth), diamonds predominantly inhibitory (net killing); conditions with no marker fall into, and contribute to, both regimes.

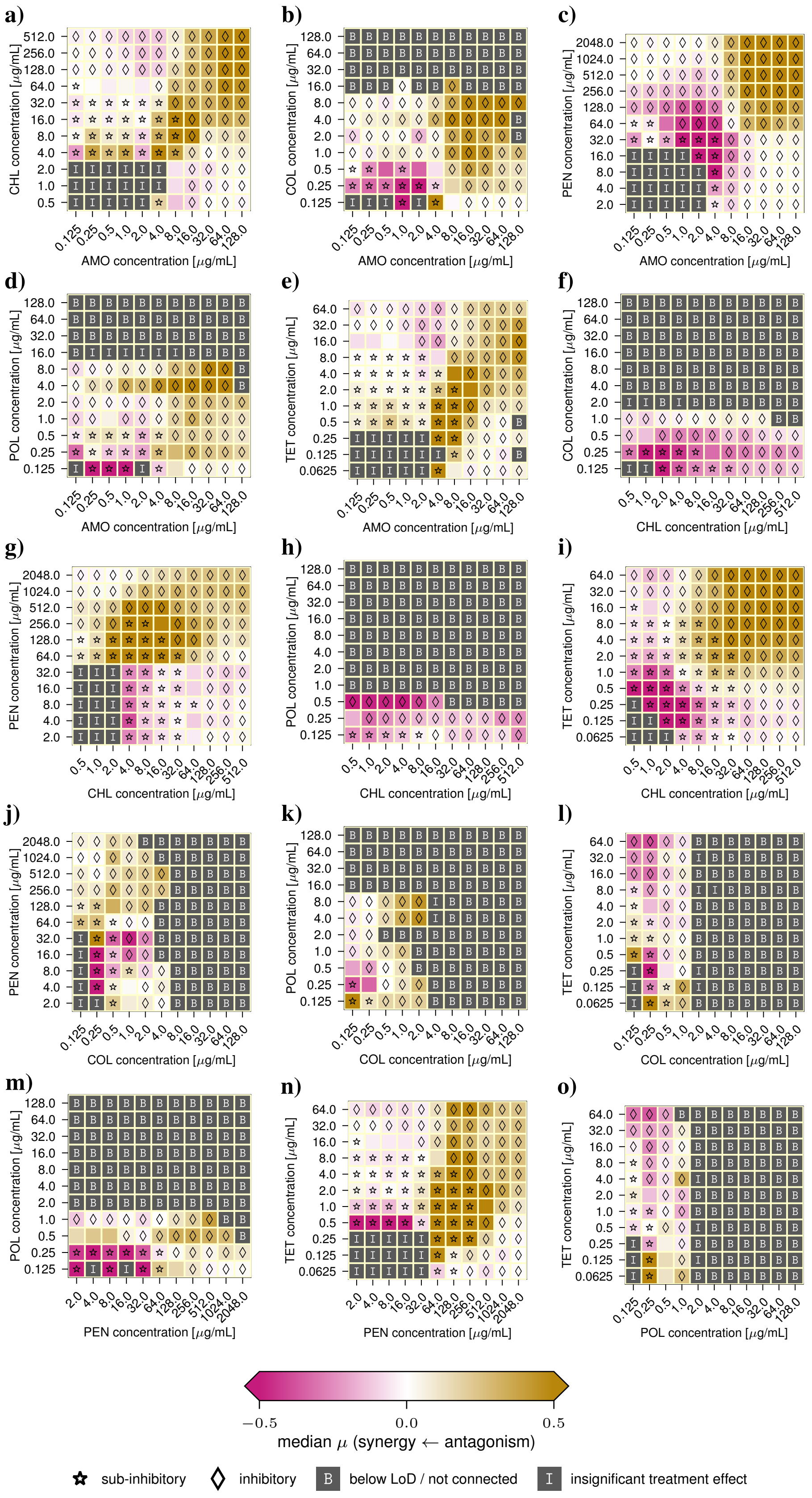


**Figure S4:** Bliss interaction ($\mu$) direction across all drug combinations. Each panel corresponds to one drug pair, and each cell represents a condition $(c_A, c_B)$. Eligible conditions are coloured by the median interaction score $\mu$ (subsection 1D): synergy ($\mu < 0$) magenta, antagonism ($\mu > 0$) gold, agreement with the Bliss prediction ($\mu \approx 0$) white. Their markers show which regime they contribute to: stars — sub-inhibitory; diamonds — inhibitory; no marker means the condition contributes to both regimes. Ineligible conditions are shown in grey and labelled by an exclusion reason: B — below the detection limit, or an isolated pocket not connected to the untreated control; I — insignificant treatment effect (see Methods).

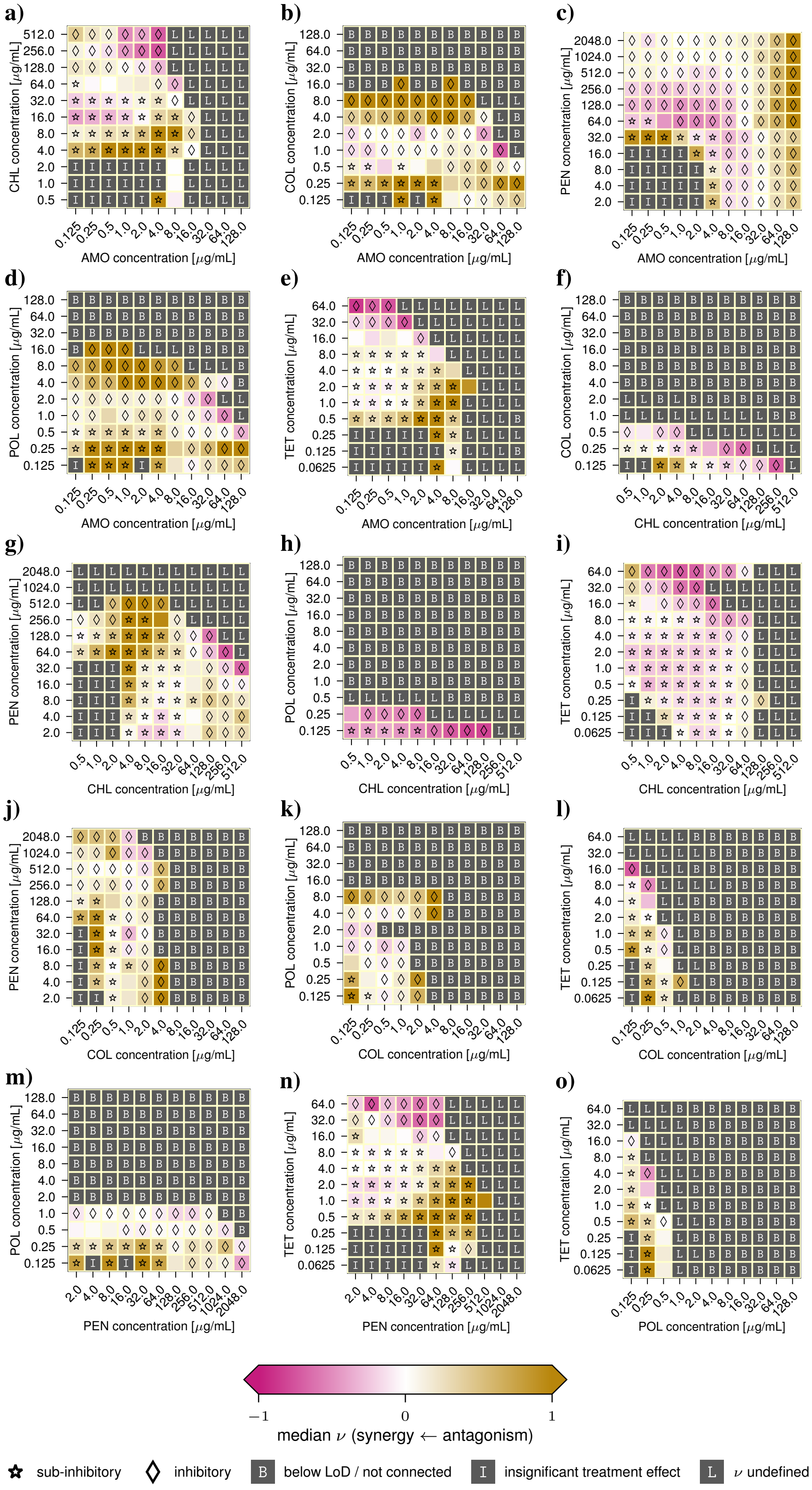

**Figure S5:** Loewe interaction ($\nu$) direction across all drug combinations. Each panel corresponds to one drug pair, and each cell represents a condition $(c_A, c_B)$. Eligible conditions are coloured by the median interaction score $\nu$ (subsection 1E): synergy ($\nu < 0$) magenta, antagonism ($\nu > 0$) gold, agreement with the Loewe prediction ($\nu \approx 0$) white. Their markers show which regime they contribute to: stars — sub-inhibitory; diamonds — inhibitory; no marker means the condition contributes to both regimes. Ineligible conditions are shown in grey and labelled by an exclusion reason: B — below the detection limit, or an isolated pocket not connected to the untreated control; I — insignificant treatment effect; L — Loewe undefined for more than 90% of the posterior draws (the combined treatment effect lies outside the effect range of at least one single drug; see Methods).

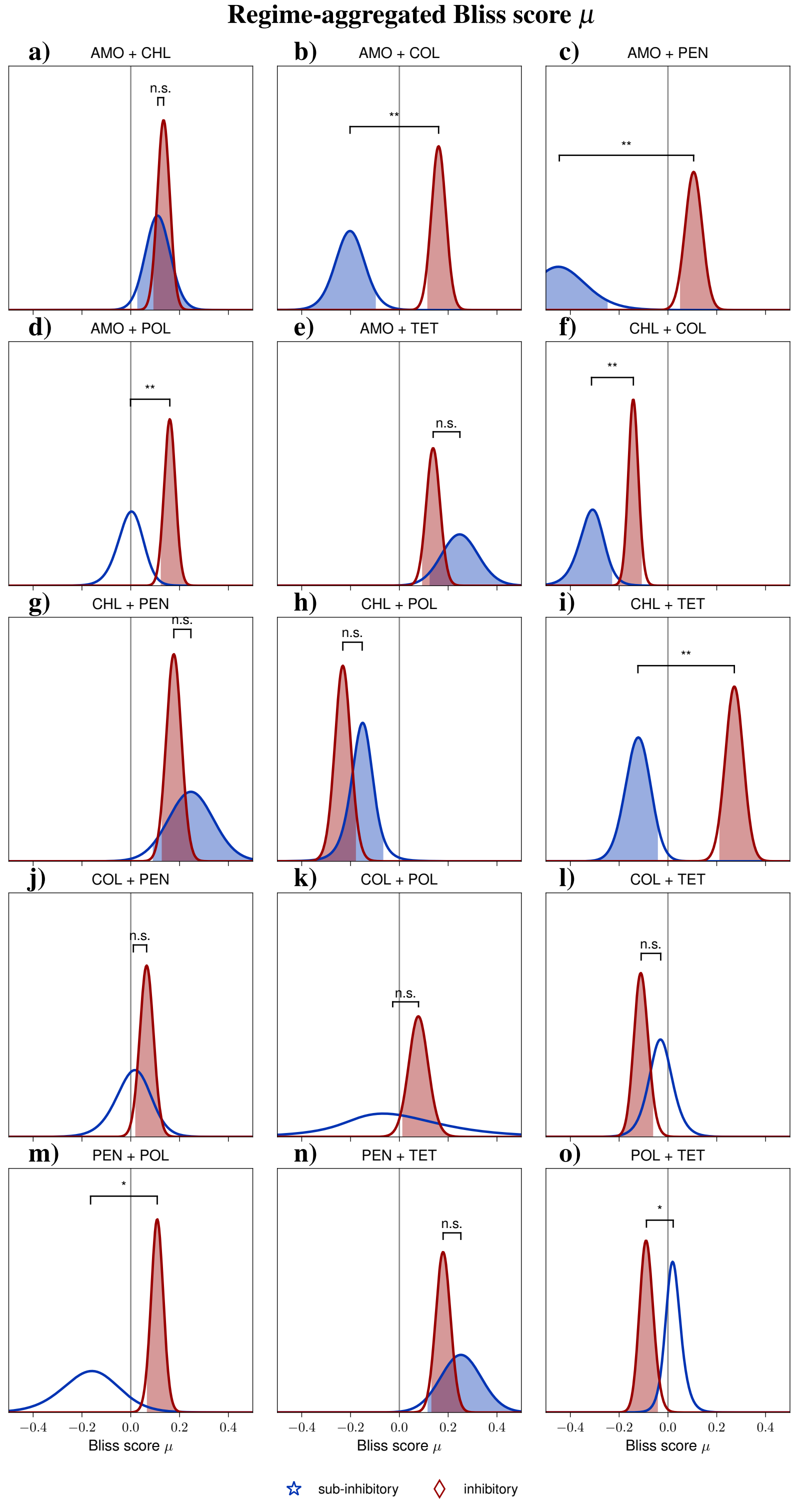


**Figure S6:** Regime-aggregated posterior density of the Bliss interaction score $\mu$ for all drug pairs (subsection 1G). The sub-inhibitory posterior is shown in blue, inhibitory in red. For regimes identified as synergistic or antagonistic, we shade the one-sided $95\%$ of the posterior mass lying on that side of zero. The grey vertical line marks zero. The black brackets connect the two regimes' medians; the annotation reports how much of the difference's posterior lies on one side of zero: $**$ at $99\%$, $*$ at $95\%$, and not significant (n.s.) otherwise.

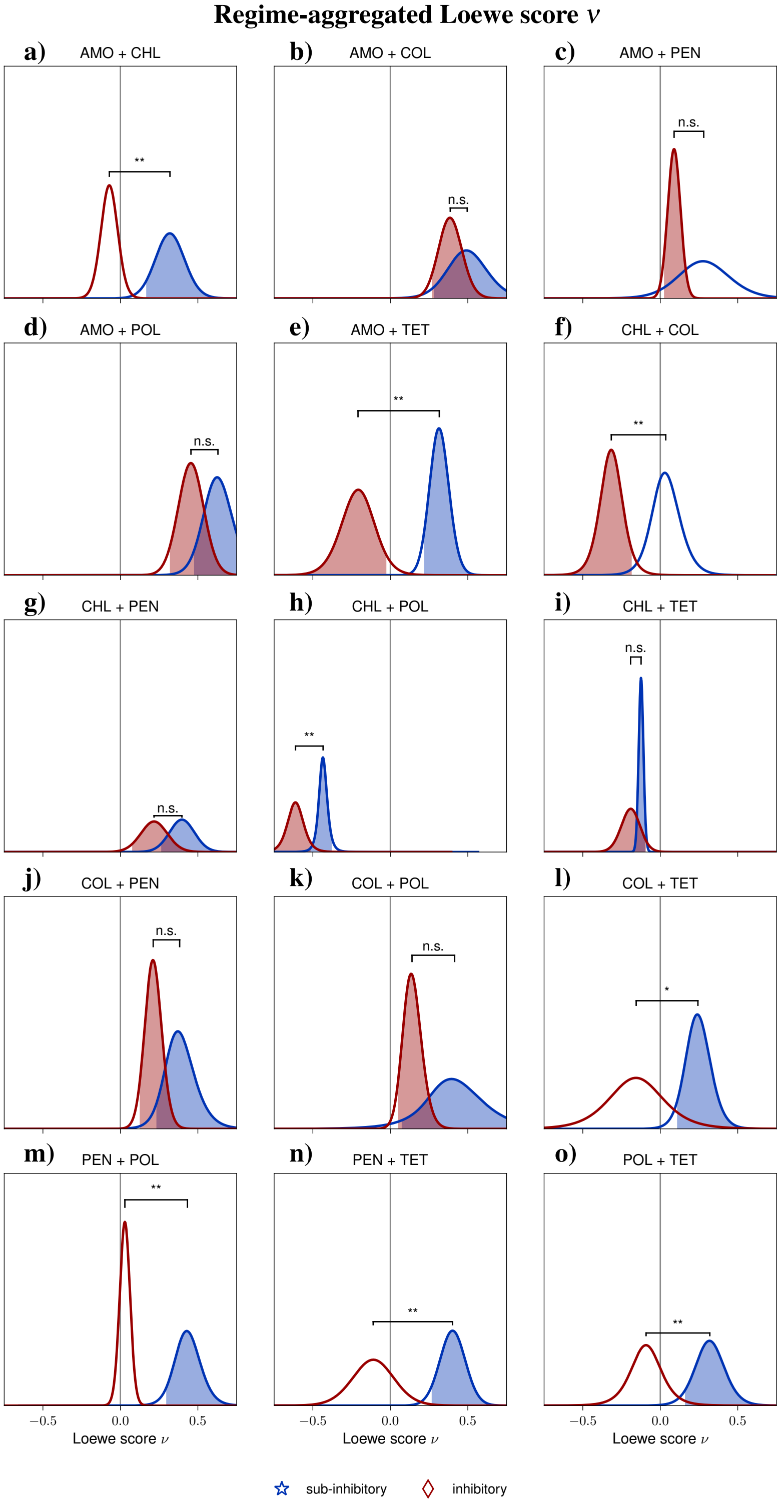


**Figure S7:** Regime-aggregated posterior density of the Loewe interaction score $\nu$ for all drug pairs (subsection 1G). The sub-inhibitory posterior is shown in blue, inhibitory in red. For regimes identified as synergistic or antagonistic, we shade the one-sided 95% of the posterior mass lying on that side of zero. The grey vertical line marks zero. The black brackets connect the two regimes' medians; the annotation reports how much of the difference's posterior lies on one side of zero: $**$ at 99%, $*$ at 95%, and not significant (n.s.) otherwise.

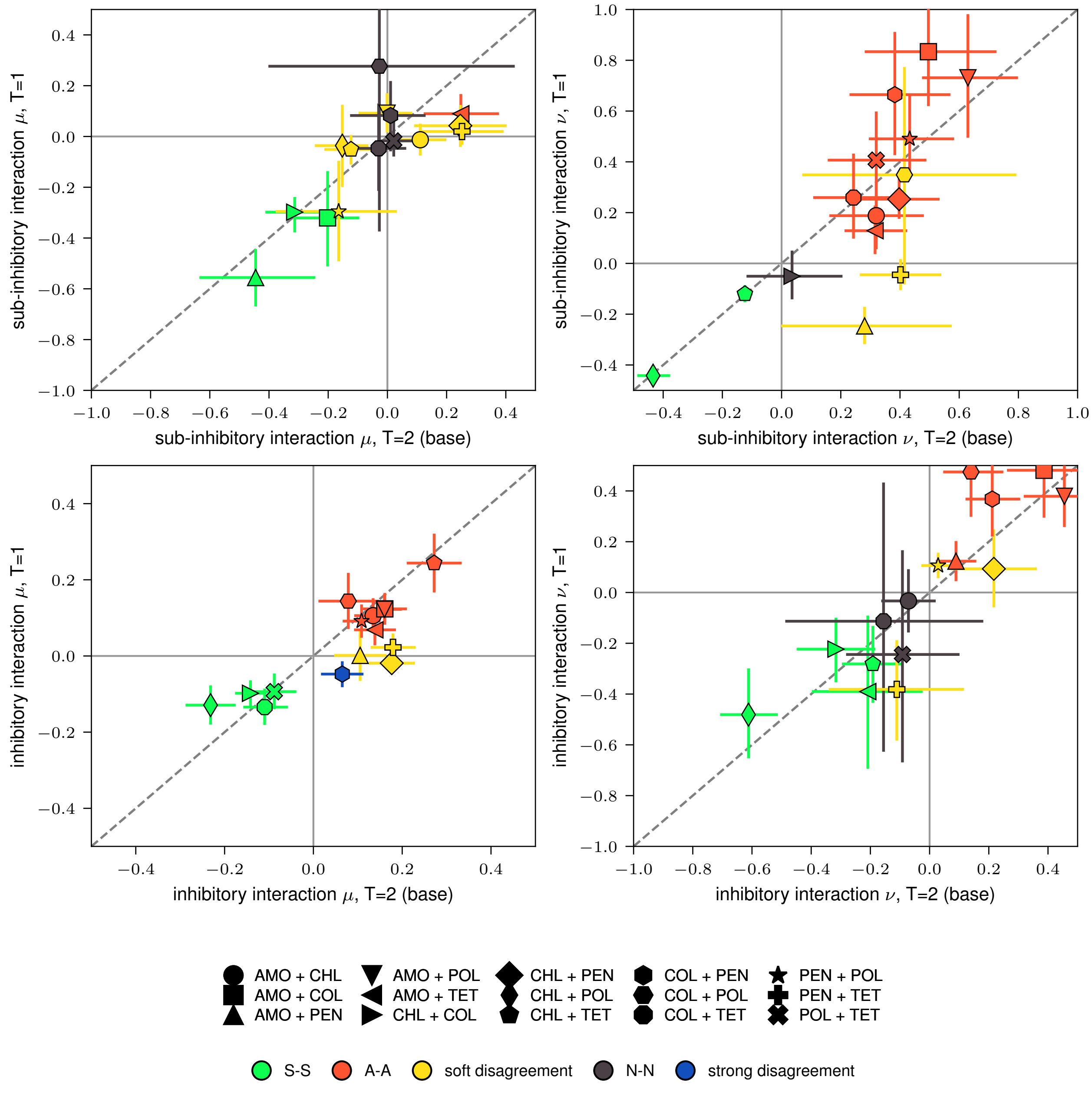


**Figure S8:** Time-window dependence of the interaction scores. Rows are the concentration regime (sub-inhibitory, inhibitory) and columns the reference model (Bliss, Loewe). Each panel scatters all drug pairs as the score for the base time window ($T = 2, x$) against the shorter window ($T = 1, y$); the dashed line marks $y = x$ (time-invariance of the posterior) and the solid axes the synergy/antagonism split. The marker encodes the drug pair and the colour the agreement of the interaction label between the two time windows for that same model and regime (i.e. whether, e.g., Loewe-inhibitory at $T = 1$ matches Loewe-inhibitory at $T = 2$). Error bars give the central $90\%$ interval of the regime posterior.

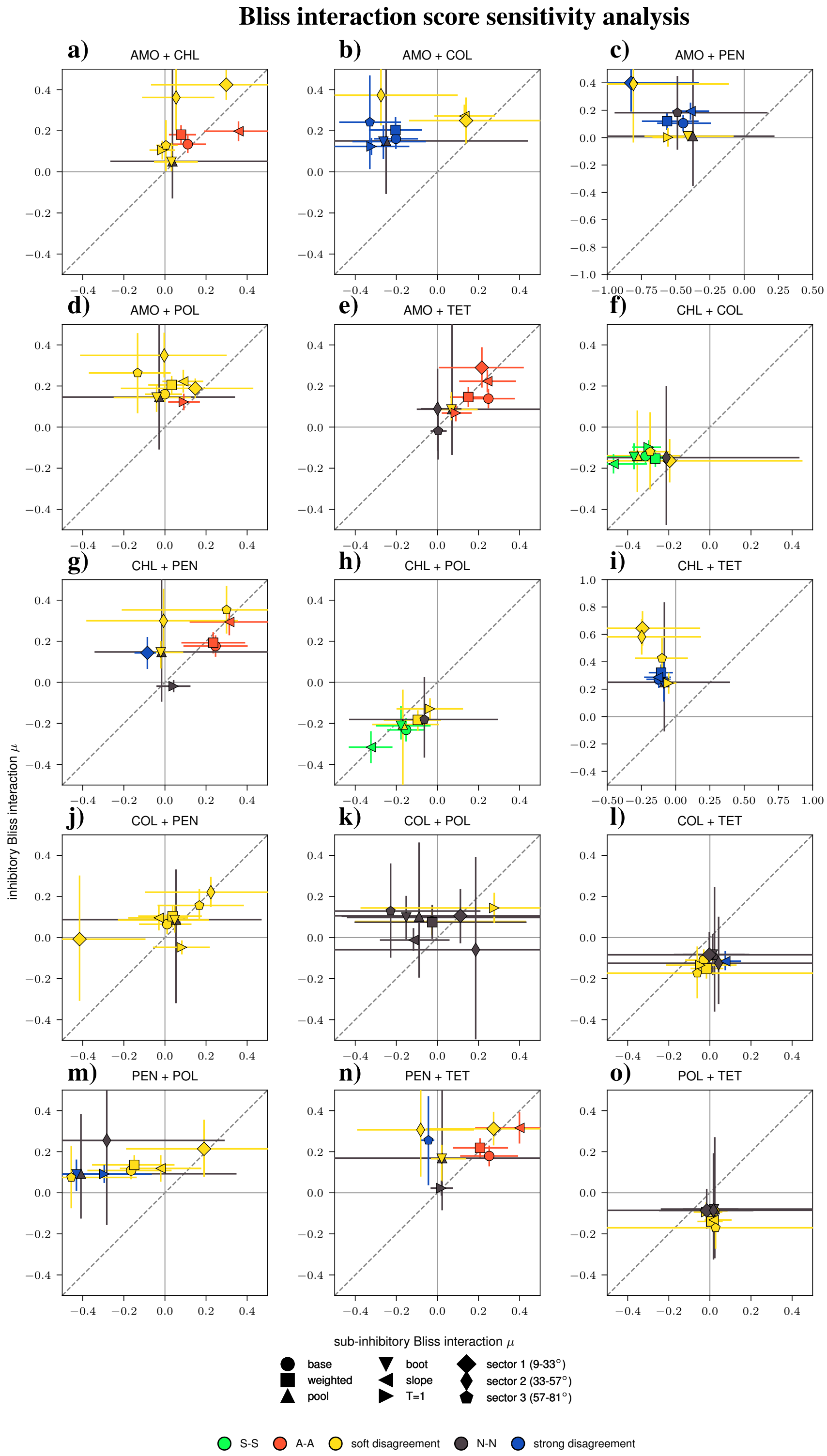


**Figure S9:** Methodological sensitivity of the Bliss interaction labels across all drug pairs. Each panel is one drug pair (titled accordingly), showing the inhibitory over the sub-inhibitory Bliss scores $\mu$ for the base analysis (time-weighted net-growth rate, $T = 2$) and eight variants: $\sin(2\phi)$-weighted aggregation, two alternative pooling schemes (pool, boot), the slope rate estimator, the shorter time window ($T = 1$), and three $\phi$-sector-restricted samples. The dashed diagonal marks $y = x$ (equal sub-inhibitory and inhibitory score), i.e. where the concentration regime does not change the score; the solid axes mark the synergy/antagonism split at zero. Error bars give the central 90% interval of the regime posterior; the marker encodes the configuration and the colour the agreement between regimes.

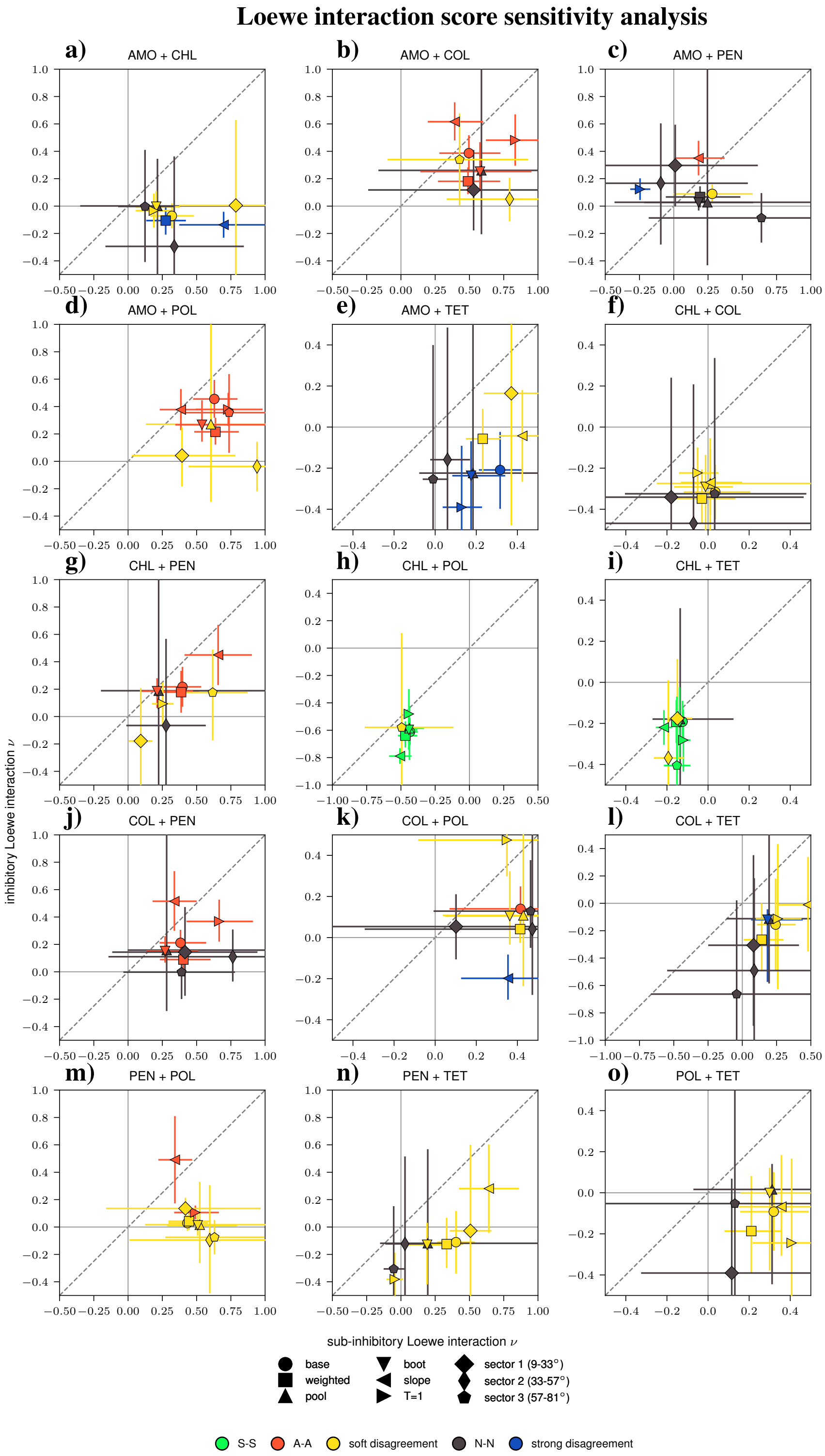


**Figure S10:** Methodological sensitivity of the Loewe interaction labels across all drug pairs. Each panel is one drug pair (titled accordingly), showing the inhibitory over the sub-inhibitory Loewe scores $\nu$ for the base analysis (time-weighted net-growth rate, $T = 2$) and eight variants: $\sin(2\phi)$-weighted aggregation, two alternative pooling schemes (pool, boot), the slope rate estimator, the shorter time window ($T = 1$), and three $\phi$-sector-restricted samples. The dashed diagonal marks $y = x$ (equal sub-inhibitory and inhibitory score), i.e. where the concentration regime does not change the score; the solid axes mark the synergy/antagonism split at zero. Error bars give the central $90\%$ interval of the regime posterior; the marker encodes the configuration and the colour the agreement between regimes.

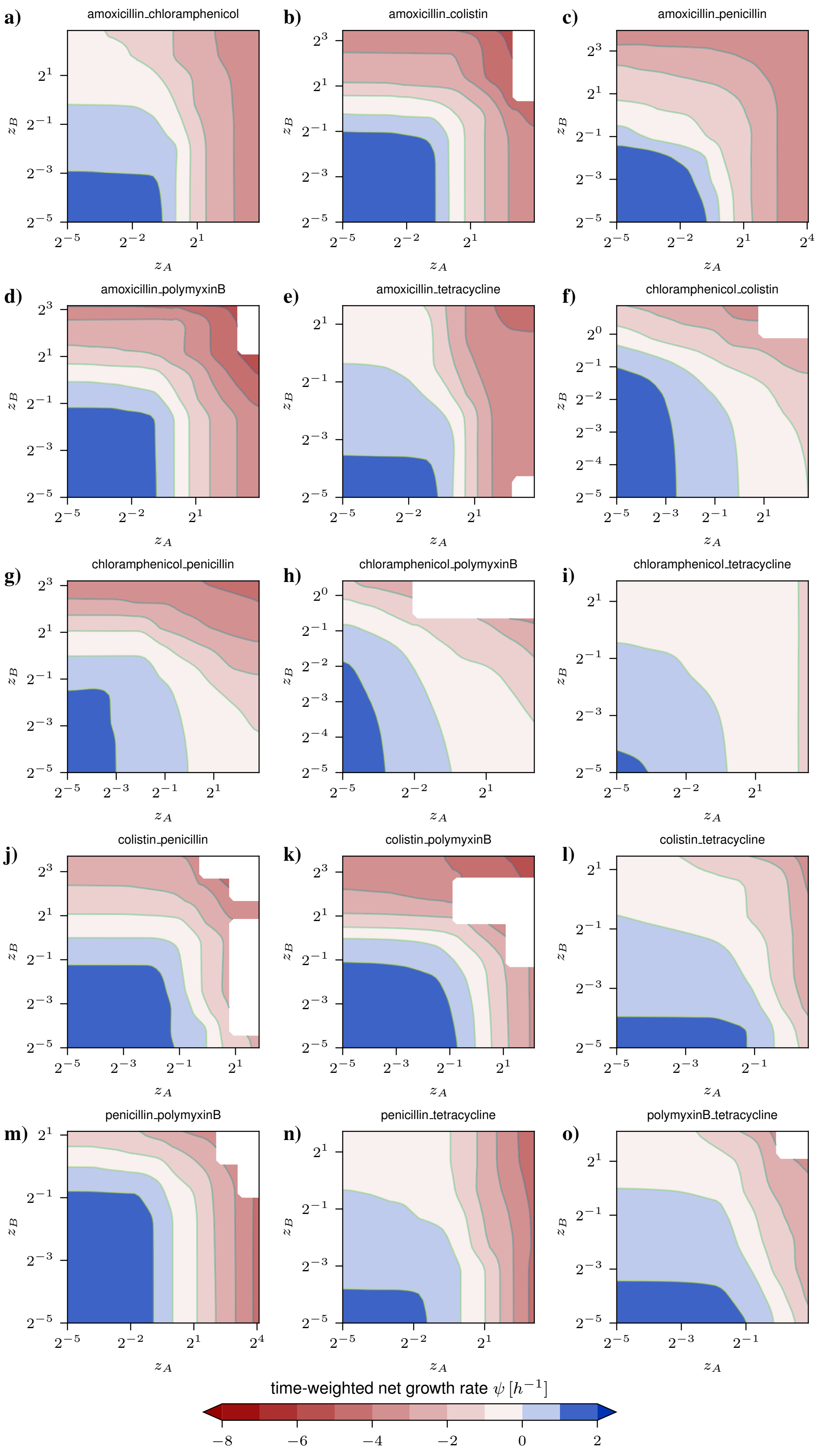


**Figure S11:** Time-weighted net growth rate surfaces for all drug combinations. Each panel shows the topography of the consensus surface (subsection 1J). Colours indicate the magnitude of $\psi$, with negative values (reds) corresponding to net killing and positive values (blues) and white regions for $\psi \approx 0$ (no net growth or killing).

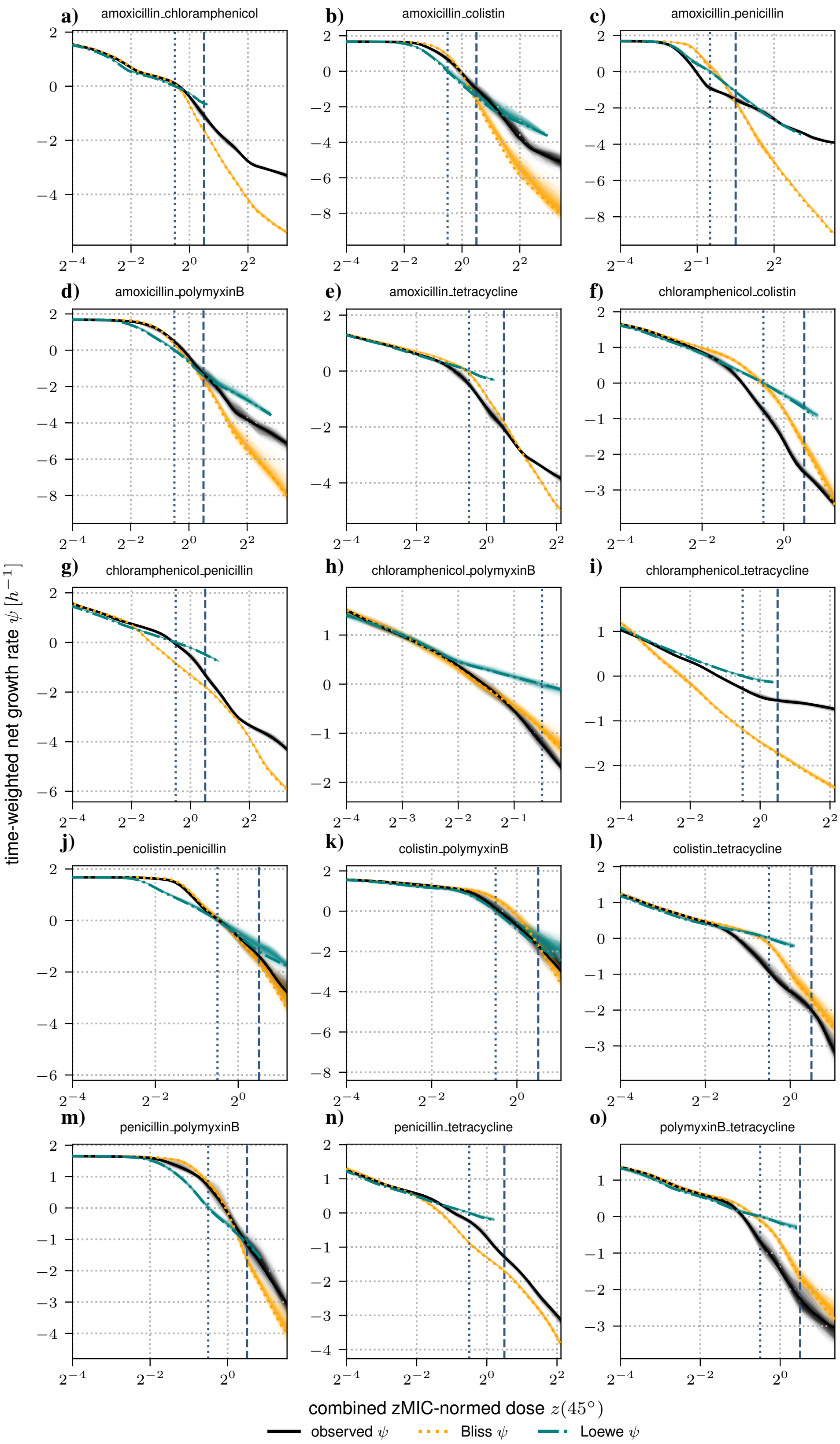


**Figure S12:** Polar pharmacodynamic curves for all drug combinations. For each drug pair, we show cross-sections through the surfaces at $\phi = 45^\circ$ (equal mixing in units of zMIC), plotting $\psi$ as a function of the combined dose $z$ (subsection 1J). The blue dotted line marks $z = 1/\sqrt{2}$, corresponding to both single-drug doses being at $0.5\,\mathrm{zMIC}$, and the blue dashed line marks $z = \sqrt{2}$, corresponding to both single-drug concentrations being at $1\,\mathrm{zMIC}$. Black, solid curves show the observed $\psi$, with corresponding Bliss (orange, dotted) and Loewe (teal, dashed-dotted) predictions evaluated along the same path $(z, \phi)$; shaded regions show the uncertainty of the surface estimate. Synergy/Antagonism corresponds to the reference model prediction of $\psi$ being above/below the observed $\psi$.

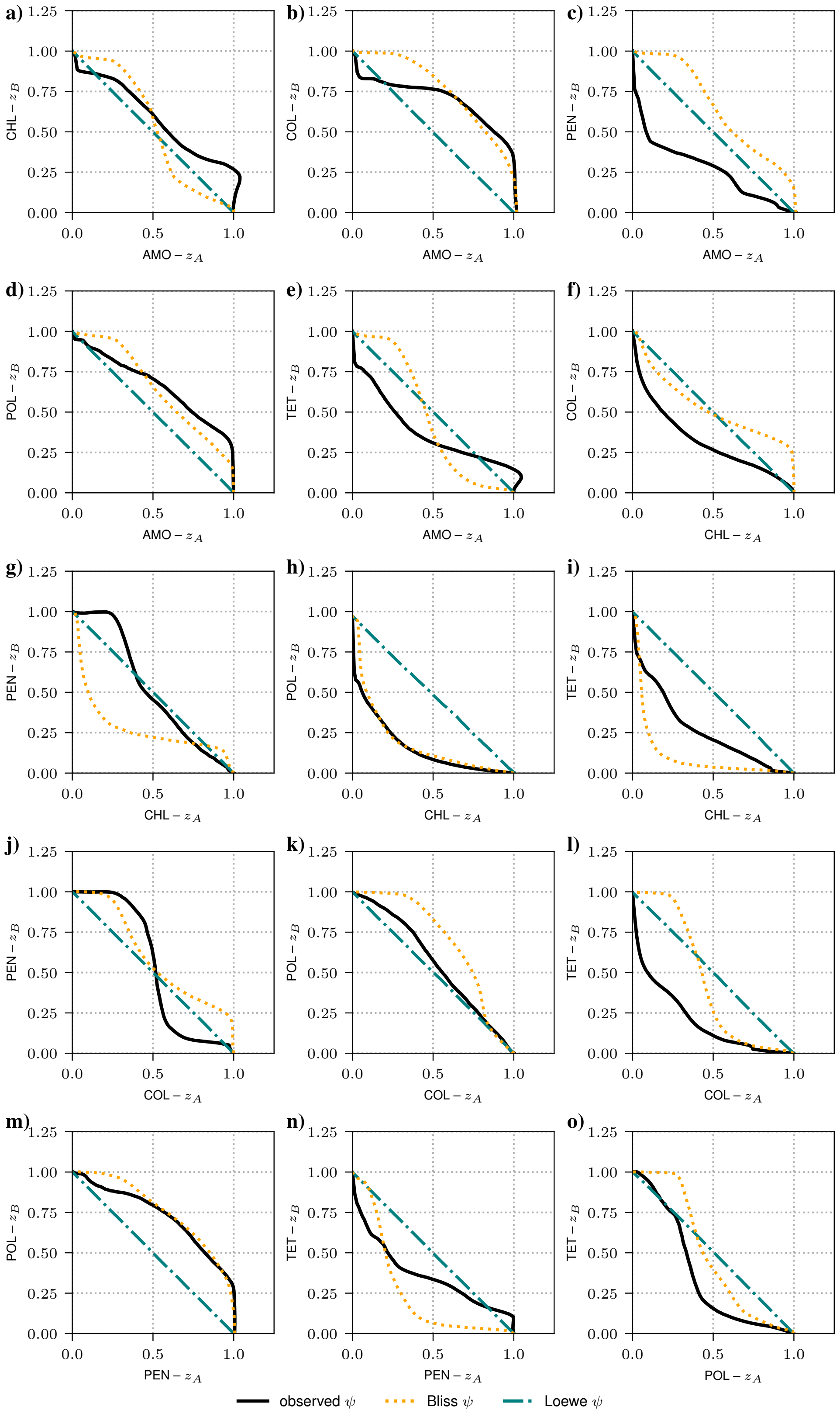


**Figure S13:** Panels a–o show isoboles of $\psi = 0\,\mathrm{h}^{-1}$ for all combinations (subsection 1J). Black solid lines show the observed isobole. In teal (dash-dotted) we plot the Loewe-based isobole and in orange (dotted) the Bliss-based isobole. An observed isobole left/below a reference model indicates synergy (less drug is needed for the same effect), right/above antagonism (more drug is needed).

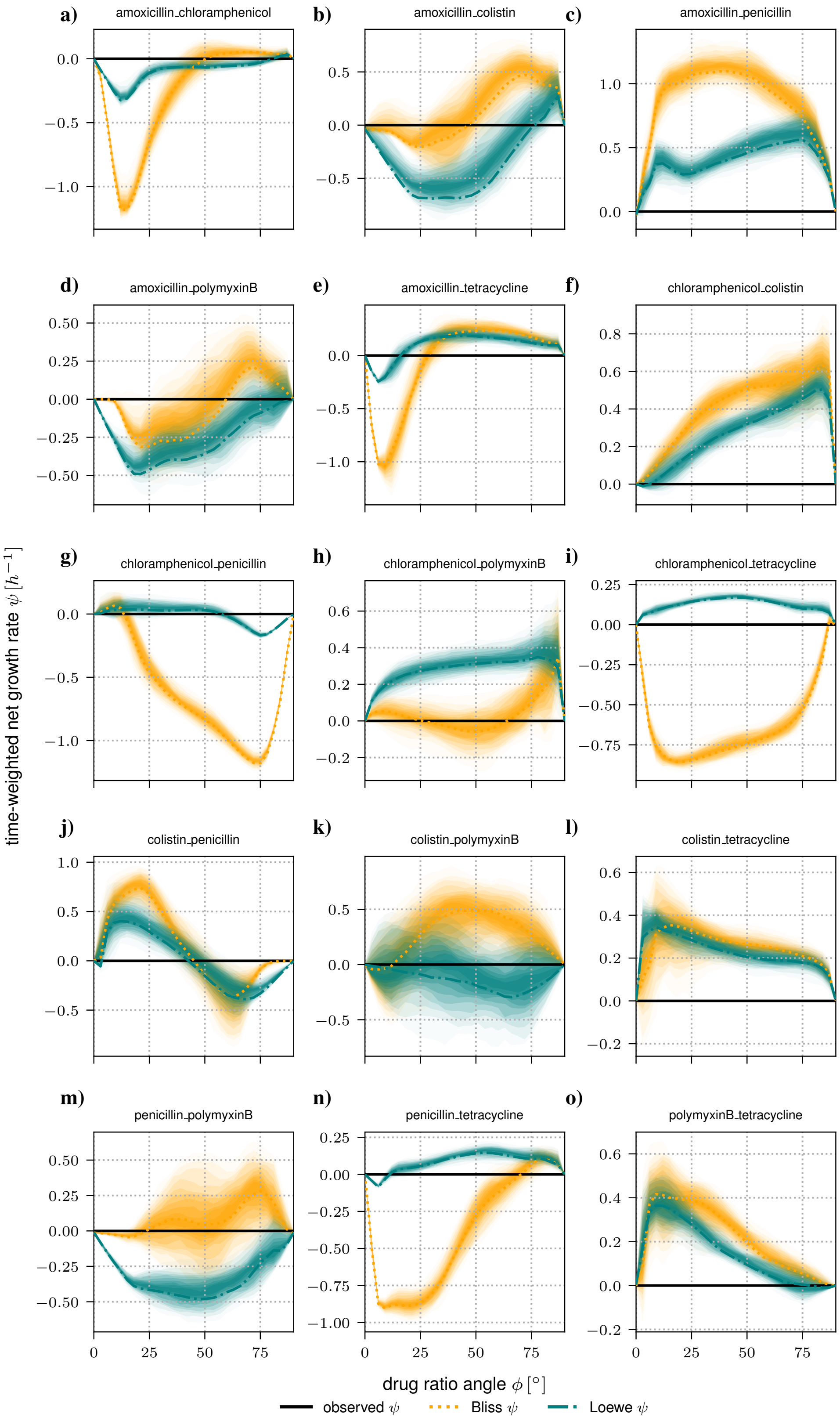


**Figure S14:** Angular interaction profiles for all drug combinations. For each drug pair, we extract isoboles (paths of constant $\psi = 0h^{-1}$) from the surfaces. Along the path $(z, \phi)_i$, we evaluate Bliss- (orange, dotted) and Loewe- (teal, dash-dotted) based predictions for $\psi$ and plot them as a function of the mixing angle $\phi$; shaded regions show the uncertainty of the surface estimate. Synergy/Antagonism corresponds to the reference model prediction of $\psi$ being above/below the observed isobole (black).

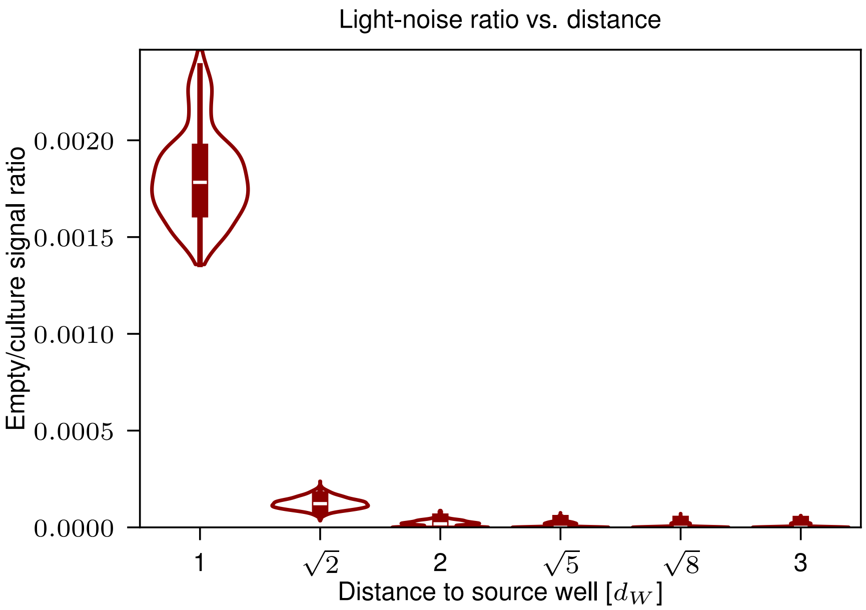


**Figure S15:** Stray light distribution, showing the fraction of light intensity emitted by a bioluminescent culture in a source well that arrives in neighbouring wells, as a function of the center-to-center distance to the source well in well lengths $d_W$. Here $d_W = 1$ corresponds to the direct horizontal or vertical neighbour, and $d_W = \sqrt{2}$ to a direct diagonal neighbour.

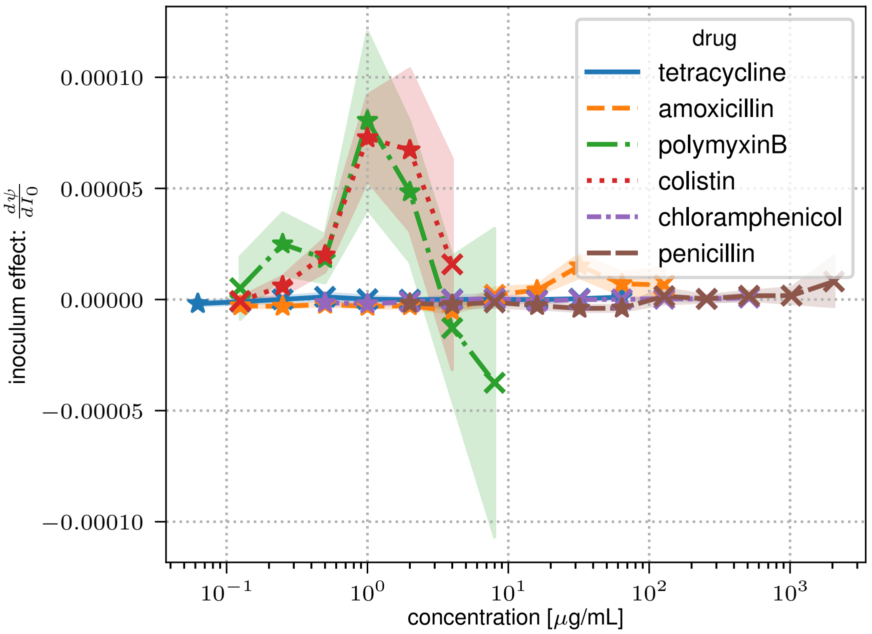


**Figure S16:** Slope of the inoculum effect ($\mathrm{d}\psi/\mathrm{d}I_0$) as a function of drug concentration. For each drug and concentration, slopes ($\mathrm{d}\psi/\mathrm{d}I_0$) are obtained by regressing the observed net growth rates across inocula, with uncertainty estimated from the regression standard error. Crosses denote concentrations with non-significant inoculum effects, whereas stars indicate statistically significant effects.
